# Integrase-On-Demand: Bioprospecting integrases for targeted genomic insertion of genetic cargo

**DOI:** 10.1101/2025.09.16.676606

**Authors:** Hannah M. McClain, Lillian C. Lowrey, Laura B. Quinto, Ellis L. Torrance, Tomas R. Gagliano, Farren J. Isaacs, Joseph S. Schoeniger, Kelly P. Williams, Catherine M. Mageeney

## Abstract

Integrases recognize defined sites within DNA and facilitate homologous recombination-independent transfer of genetic fragments. These innate capabilities render integrases as powerful biotechnology tools that mediate efficient site-specific integration of large genetic cargos. Given that genomes often lack the integration sites recognized by frequently utilized integrases, integrase technology has largely been restricted to genetic engineering of model organisms into which *attB* sites can be synthetically introduced. To enable single-step site-specific integrase-mediated genome editing in a broad spectrum of prokaryotes, we have devised the *Integrase-On-Demand* (IOD) method. IOD systematically identifies integrases, from bacteria and archaea, that can integrate into available *attB* sites in any target prokaryote. Computational results show that diverse bacteria generally have multiple potentially useable native *attB*/integrase pairs. We confirmed the functionality of predicted integrase and *attB* pairs for mediating site-specific genomic integration of heterologous DNA into the genomes of *Pseudomonas putida* S12 and KT2440 and *Synechococcus elongatus* UTEX 2973 and measured efficiency of integration using suicide constructs. By eliminating the requirement to introduce non-native *attB* sites into the target genome, IOD may, when suitable transformation methods exist, allow facile genome integration of large constructs in non-model and possibly non-culturable bacteria.

**Graphical Abstract:** 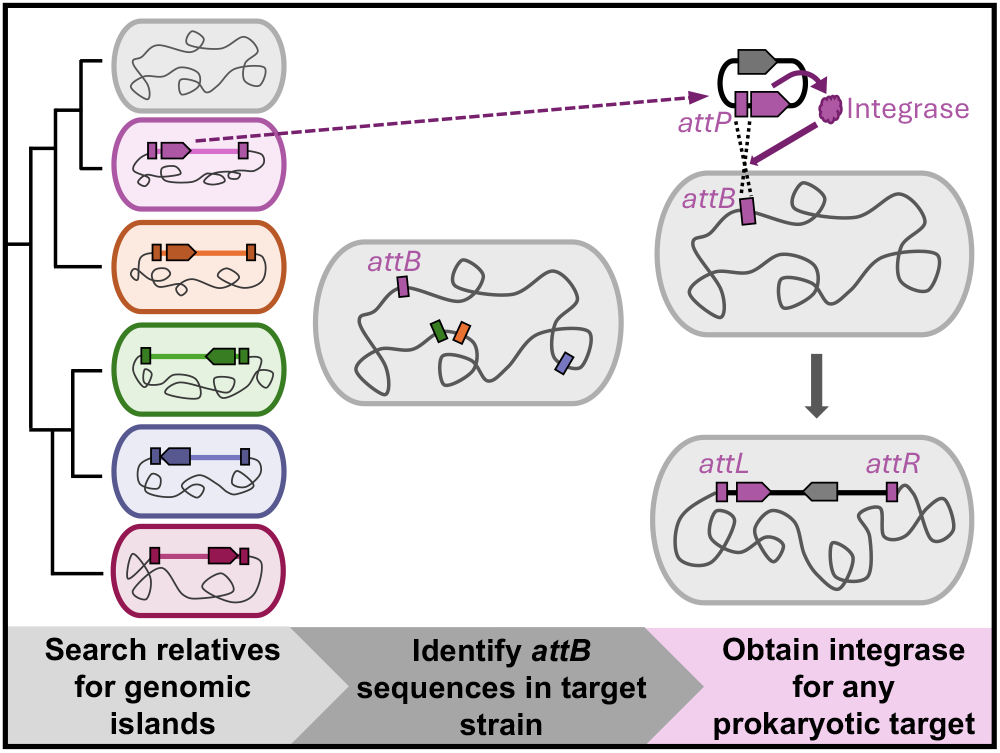

## INTRODUCTION

Bacterial genome engineering enables basic studies of genetics and functional genomics of bacteria and microbiomes and is driving advances in biotechnology applications such as biomanufacturing (1), bioremediation (2), soil health and rhizosphere adaptation (3), and human health (4). As biotechnology applications have diversified, and research into microbial ecosystem studies enabled by metagenomics has expanded, less well-studied, genetically and metabolically diverse microbes have elicited interest. These often lack established genetic tools, or may be difficult to isolate or culture, which impedes the use of genetic tools for research and synthetic biology efforts to introduce genes conferring novel or enhanced functions.

Integrases, site-specific DNA recombinases, provide a potential solution for efficiently introducing large genetic cassettes into non-model and possibly unculturable bacteria (5,6). Integrases have many advantages over other methods of genetic engineering including insertion of large DNA payloads (6), scarless insertions (7), conservation of *attB* sites between related taxa (8), and site-specific insertion with minimal off-target activity (9). Technologies have been developed to utilize integrases for genome engineering (10), including programmable addition via site-specific targeting elements (PASTE) (11), recombinase mediated cassette exchange (RMCE) (12) and serine recombinase-assisted genome engineering (SAGE) (13). However, these methods first require the introduction of non-native *attB* sites as a ‘landing pad’ (i.e. integration site) for subsequent DNA cargo integration. Landing pads are typically introduced using transposons, which are not site-specific. Additionally, this two-step transformation model compounds inefficiencies in each step, and hinders widespread adoption of integrase technologies, especially in organisms for which transformation is difficult or there are no robust genetic engineering tools. Newer integration methods with “programmable” integration site-specificity such as cas-transposon systems (14) and bridge RNA-guided recombination (15) promise to have great utility, but suffer, compared to classical integrases, from limited efficiency, host range and insert size, for the former, and higher potential for off-target site integration or mis-sequencing if multiple integration steps are performed, for the latter.

Many prokaryotic genomes harbor integrases within mobile integrative genetic elements (hereafter referred to as genomic islands (GIs)). GIs are discrete regions of chromosomal DNA that contain an integrase gene, typically differ in sequence composition from the host genome, and may be shared sporadically among closely related bacteria (16). The GI-encoded integrase enzyme facilitates integration of a mobile genetic element (MGE) (e.g. a temperate phage or integrative conjugative element) into the host chromosome to form the GI by catalyzing DNA recombination between attachment sites (*att*) in the MGE (*attP*) and in the genome (*attB*) (7). Recombination produces two hybrid *att* sites, *attL* and *attR*, which flank the GI within the chromosome and serve as substrates for a potential subsequent GI excision reaction that requires both the integrase and a Recombination Directionality Factor (RDF) or excisionase (*xis*) gene (17). Two unrelated integrase families exist, tyrosine and serine recombinases, and while serine integrases are more commonly utilized in biotechnology (5), both families are capable of catalyzing integration, inversion, and excision of precisely defined regions of DNA.

To facilitate site-specific genomic insertion of large genetic cargo in a wide range of prokaryotes (both bacteria and archaea), we have developed the Integrase-On-Demand (IOD) method. Recent advances in precise GI discovery, enabled by our Islander (18) and TIGER (16,19) software, have greatly expanded the discovery of integrases and precisely predicted *att* sites for both tyrosine and serine integrase GIs across hundreds of thousands of diverse prokaryotic genomes (20). IOD starts with a computational pipeline that leverages these advances to identify prokaryotic serine and tyrosine integrases predicted to recognize *attB* sequences present within a user-defined bacterial or archaeal strain of interest. Here we demonstrate that the IOD software can predict integrases and *att* sequences from diverse bacteria and archaea. As a proof-of-concept for the IOD method, we experimentally validated that suicide vectors could site-specifically, efficiently, and stably insert DNA constructs into the chromosomes of two phylogenetically diverse and biotechnologically relevant bacterial species: the gammaproteobacterium *Pseudomonas putida* (strains S12 (nine integrases) and KT2440 (three integrases)), and the cyanobacterium *Synechococcus elongatus* UTEX 2973 (one integrase).

## MATERIALS AND METHODS

### Development of IOD software

IOD references a list of *attB* sequences and their corresponding integrase proteins to provide the user with a list of candidates. Each *attB*/integrase pair is derived from a GI predicted by the TIGER/Islander programs (16,18,19). The programs have been run on over 450,000 genomes, yielding 1,757,053 GIs containing integrases paired with their cognate *att* sites (20). To build the reference data structure, the set of unique *attB* sequences of the predicted islands were collected.

Integrases theorized to be inactive due to truncation or lack of catalytic domains were excluded from the analysis. To assess tyrosine integrase integrity, we compared these integrases to the 20 HMMs of the tyrosine recombinase database (TRDB) (21). Putative tyrosine integrases with no HMM match, or that matched to the Xer or Integron HMMs (subfamilies that perform non-integrase functions) were rejected. Furthermore, tyrosine integrases longer than 800 amino acids were rejected. Finally, a metric (ΔΔHA) was developed to identify truncated proteins, calculating the number of amino acid positions of the HMM missing at the termini of the protein (ΔH) and the number of amino acids at the termini of the protein exceeding the HMM hit (ΔA), and taking ΔΔHA = ΔH - ΔA. We used a cutoff of ΔΔHA>100, which passed 98.5% of the medium-sized tyrosine integrases (250-800 aa) and 8.9% of the short tyrosine integrases (<250 aa), to reject nominally truncated proteins. Lastly, serine integrases were rejected unless: 1) it had the Pfam Resolvase domain followed by the Pfam Recombinase domain and was not too long (>800 amino acids) or too short (ΔΔHA > 0); or 2) our Tater software (16) had previously classified it as S-Core_IS607 and it was not too long (>300 aa) or too short (ΔΔHA > 0). As a result, the final list of *attB* sites queried by IOD is 182,734 unique sequences (**Figure 1**).

**Figure 1.**
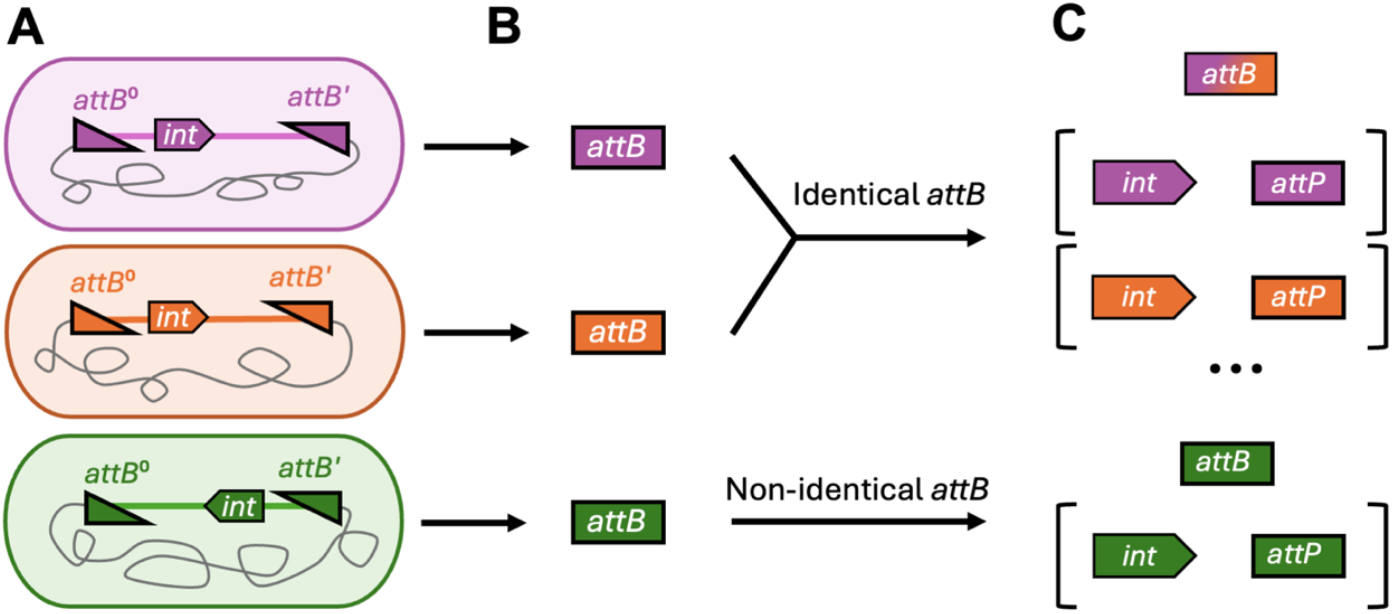
Preprocessing steps for IOD Integrase and *att* pairs. **A)** *attB* sites are extrapolated from TIGER/Islander predicted genomic islands. **B)** *attB*-integrase pairs are built based on *attB* sequence identity. **C)** At least one integrase and predicted GI is associated with each *attB* sequence.

The program features two modes (*search* and t*axonomic*) along with several command line options. These arguments are handled in the wrapper script *runiod*.*py. Search* mode allows the user to query any sequence against the IOD database for *attB* sequences. If a match is found in the genome of interest, the *attB* site, as well as the sequence(s) of the corresponding integrase(s), are returned to the user. *Search* mode can be invoked by adding “-*search”* as a flag. The user may provide specific sequences to search by using “*-seq*” followed by one or more comma separated sequence(s) directly to the command line or the path to a file containing sequences in either raw or FASTA format. If no sequences are provided in *search* mode, then the whole database is used. Users have options to modify the minimum number of *attB* sequences to collect in *taxonomic* mode prior to searching (default: n=500), length of *attB* flanks paired to serine (default: sl=16), and tyrosine (default: yl=10) integrases, and number of threads for BLAST to use (default: threads=1).

*Taxonomic* mode runs MASH (22), a program that uses a hash function to reduce genomic sequences to representative sketches. A sketch of the user-provided genome is created and compared to a pre-compiled database of 85,205 GTDB representative species (23), returning a list of genomes which have high similarity to the query. Starting with the best candidate species, the *attB* sequences and their respective integrase(s) from the IOD database corresponding to the queried species, are collected until the arbitrarily defined minimum number (default: n=500) are retrieved. If the minimum number is not met, then pairs from the next most closely related species are collected. If the sum of all pairs for all candidate species is still less than the minimum number of pairs specified by the user, then the reference *attBs* are collected from all members of the species’ genus until the numerical quota is met. Once the numerical quota is met, a query FASTA including each *attB, attB’* (left flank and identity block of *attB*), and *attB*^*o*^ (identity block and right flank of *attB*) is written to the output directory (the command line arguments *-yl* and *-sl* can be used to modify the flank length). The half *attB* sites, *attB’*, and *attB*^*o*^ are intended to probe the genome of interest for GIs in the database. Next, a nucleotide BLAST database is produced from the user-provided genome (24). The *attBs* are searched against the BLAST database of the genome using *-task blastn*. The hits with >95% identity, one or no gaps in the alignment, and a maximum of two base pair mismatches are categorized as matching to the database *attB, attB’*, and *attB*^*o*^. The categorized *attB* blast hits are then filtered to determine if the site is a candidate or not. If the reference *attB* is determined to have a single *attB h*it in the query, then it passes through the first candidate filter. Frequently, a sequence may have more than one full or partial *attB* hit, these hits do not pass the candidate filter as they may have an island occupying the prospective *attB* site.

For *attB* sites removed from the candidates list, the length of each reference island is collected to determine if and which island is occupying the integration site. Each putative island length is then compared to the distance between the *attB’* and *attB*^*o*^ hits. The program can also handle *attBs* that span contig boundaries. If the hits map to different contigs, then the direction of the BLAST hit is used to determine which contig termini would be expected to be part of a GI. To be predicted as a genomic island, the normalized (for length of expected island) difference must be less than 0.05:

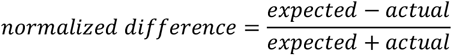

The candidate list is then passed through another series of filters. These check the candidate IOD matches for questionable features such as low-support reference islands, multiple copies of the ID-block (estimated crossover site), and tandem repeats in the ID-block.

Despite deduplication of sequences in pre-processing, the list of candidates often contains overlapping *attB* sites. The shifted *attB* sites could be explained by mutations between closely related species that share a GI and/or the limit of precision in integration sites called by TIGER/Islander software (16,18,19). The program bins all candidates by overlapping coordinates and attempts to select the highest quality sequence from each bin according to the best BLAST support score. If there are ties, then the site whose reference island has the highest support will be selected. The best candidate *attBs* are reported in *final_candidates*.*tsv*. An example reference island is reported according to the highest support, along with pairs of integrases and the island ID (GCA accession and gene ID). All *attB* sites are reported with each associated reference island in the isles.*json* files.

Once genomic sites inferred to contain an island are deduplicated by overlapping coordinates, they are reported in *occupied*.*tsv* along with the reference genomic islands. Occasionally, candidate *attBs* are found *within* occupied islands, and these are reported in the *attb_dupes* files with the island in which they are contained.

### Software Testing

Two sets of genomes were collected for study. The first set (Health) contained 41 genomes of health and biomanufacturing relevance. The second set (Diversity) contained 142 genomes each representing a different GTDB (23) phylum with more than 4 genomes available.

Both genome sets were tested in *taxonomic* and *search* modes using default parameters (**Table 1 and Supplementary Table 1**). *Taxonomic* mode searches the query genome for up to 500 *attB* sequences from its close relatives, whereas search mode uses 182,734 *attB* sites. To simulate a typical user experience, the software was tested locally on a MacBook Air, M15,13. IOD was written in python (v3.12) and requires the external programs MASH (v2.3) (https://github.com/marbl/Mash) and the BLAST search algorithm to be available as a system-wide executable. The program can be run from any command line interface. Memory consumption was found to peak at 14.6MB and 14.4MB, for *taxonomic* and *search* mode respectively (**Supplementary Figure 1**). The software is available at https://github.com/sandialabs/Integrase-On-Demand.

**Table 1.** Taxonomic mode IOD output summary for the Health genome set of 41 organisms relevant for human health or biomanufacturing. GTDB species name, the NCBI genome assembly accession number, number of *attB* sequences used, number of *attB* candidates, subset of *attB* candidates sourced from islands targeting tRNA genes, and number of occupied sites are listed. **Supplementary Table 1** has the full data set with search mode results included.

| GTDB species | GCA | *attB* queries | *attB* candidates | *attBs* in tRNA genes | Occupied |
| --- | --- | --- | --- | --- | --- |
| *Bacillus coagulans* MA-13 | 004359975 | 500 | 3 | 2 | 0 |
| *Corynebacterium glutamicum* ATCC13032 | 000011325 | 152 | 10 | 7 | 2 |
| *Bacteroides fragilis* | 000025985 | 500 | 38 | 12 | 10 |
| *Corynebacterium diphtheriae* | 001457455 | 203 | 16 | 7 | 3 |
| *Acinetobacter baumannii* | 009759685 | 500 | 34 | 10 | 8 |
| *Clostridioides difficile* DSM28196 | 003482035 | 500 | 28 | 0 | 28 |
| *Clostridium tyrobutyricum* ATCC25755 | 000359585 | 286 | 8 | 3 | 9 |
| *Pseudomonas aeruginosa* | 001457615 | 500 | 46 | 15 | 6 |
| *Zymomonas mobilis* ZM4 | 003054575 | 111 | 0 | 0 | 0 |
| *Pseudomonas putida* S12 | 000495455 | 500 | 37 | 15 | 7 |
| *Clostridium ljungdahlii* DSM13528 | 000143685 | 214 | 1 | 0 | 5 |
| *Sulfolobus acidocaldarius* | 000012285 | 33 | 0 | 0 | 0 |
| *Streptomyces venezualae* | 008639165 | 500 | 20 | 10 | 16 |
| *Rhodobacter sphaeroides* ATCC17023-2.4.1 | 000012905 | 500 | 9 | 6 | 10 |
| *Pseudomonas putida* KT2440 | 000007565 | 500 | 32 | 12 | 11 |
| *Novosphingobium aromaticivorans* DSM12444 | 000013325 | 500 | 10 | 8 | 4 |
| *Haloferax volcanii* | 000025685 | 500 | 10 | 5 | 12 |
| *Clostridium acetobutylicum* ATCC824 | 000008765 | 500 | 0 | 0 | 1 |
| *Vibrio cholerae* | 000621645 | 500 | 27 | 10 | 3 |
| *Streptomyces griseus* ATCC13273 | 003610995 | 500 | 30 | 15 | 7 |
| *Methylobacterium aquaticum* | 001043915 | 500 | 6 | 6 | 5 |
| *Synechococcus elongatus* | 000817325 | 72 | 1 | 1 | 0 |
| *Methylobacterium extorquens* | 900234795 | 500 | 23 | 20 | 6 |
| *Escherichia coli* | 003697165 | 500 | 44 | 8 | 8 |
| *Enterococcus faecalis* | 000392875 | 500 | 38 | 5 | 11 |
| *Klebsiella pneumoniae* | 000742135 | 500 | 37 | 9 | 11 |
| *Bacillus subtilis* | 000009045 | 500 | 16 | 2 | 3 |
| *Bacillus licheniformis*<br>DSM13 | 000011645 | 500 | 24 | 9 | 2 |
| *Helicobacter pylori* | 900478295 | 500 | 22 | 1 | 0 |
| *Bacillus coagulans*<br>DSM2314 | 006716385 | 482 | 4 | 2 | 0 |
| *Neisseria gonorrhoeae* | 003315235 | 500 | 17 | 12 | 10 |
| *Pseudomonas fluorescens*<br>SBW25 | 000009225 | 500 | 29 | 16 | 10 |
| *Listeria monocytogenes* | 900187225 | 500 | 38 | 7 | 0 |
| *Cupriavidus necator* H16 | 004798725 | 500 | 27 | 17 | 5 |
| *Acidithiobacillus*<br>*ferrooxidans* | 000021485 | 293 | 24 | 18 | 2 |
| *Streptococcus*<br>*pneumoniae* | 001457635 | 500 | 43 | 3 | 15 |
| *Leptospirillum ferriphilum* | 000755505 | 67 | 11 | 11 | 3 |
| *Staphylococcus aureus* | 001027105 | 500 | 25 | 2 | 6 |
| *Salmonella enterica* | 000006945 | 500 | 34 | 9 | 9 |
| *Mycobacterium*<br>*tuberculosis* | 000195955 | 500 | 24 | 16 | 5 |
| *Francisella tularensis* | 000008985 | 500 | 3 | 2 | 0 |

### Phylogenetic Analysis

The protein sequence for each integrase predicted by the IOD software was obtained using the integrase coordinates. Serine and tyrosine integrase sequences were treated separately. Protein sequences were aligned using MAFFT v7.526 (25) (mafft --auto) and maximum-likelihood trees were prepared using RAxML-NG v1.2.2 (26) (raxml-ng --all --msa tyr.afa --model LG+G8+F –tree pars 10 --bs-trees 200). The resultant phylogenetic tree was viewed using FigTree v1.4.4 (http://tree.bio.ed.ac.uk/software/figtree/) and rooted at the midpoint.

### Protein alignments for catalytic residue confirmation

We used the same method as described above to create the aligned protein sequence separately for the serine integrases and tyrosine integrases. For the serine integrases we obtained the integrase reference sequences from Uniprot (27) using the following accession numbers: PhiC31 (Q9T221) and Bxb1 (Q9B086). We obtained Lambda (P03700) and Cre (P06956) as model tyrosine integrases. The aligned protein sequences were loaded into the SnapGene viewer to create the final figure and confirm catalytic residue positions.

### Genomic island prediction, induction, and recombination measurements

TIGER (16,19) and Islander (18) were used to predict GIs in *P. putida* S12 (assembly: GCF_000495455.2).

*P. putida* S12 was grown overnight at 30ºC shaking in LB broth, diluted 3:100, and grown to an OD_600_ of 0.4 and 1 ug/mL of mitomycin C (MMC) was added. Samples (1 mL) were collected at 0, 1, 2, and 3 hours post MMC addition. Cells were pelleted at 16,000 × *g* for 2 min and genomic DNA was isolated using the Qiagen DNeasy blood and tissue kit (Qiagen, Venlo, Netherlands, 69506). Genomic DNA libraries were prepared using the Illumina DNA prep kit (Illumina, San Diego, CA, 20060060; Indexes:Illumina, San Diego, CA, 20018708). Libraries were pooled in eqi-molar fashion, and the final library was sequenced using an Illumina NextSeq 500 sequencer with the high-output 300-bp single-end read sequencing kit (Illumina, Sand Diego, CA, 20024908).

Sequencing reads were quality filtered using BBDuk (v36.11; http://jgi.doe.gov/data-and-tools/bb-tools/) with the following parameters specified: ktrim=r, k=21, mink=11, hdist=1. The filtered reads were analyzed with Juxtaposer (28) to identify non-canonical reads compared to the *P. putida* S12 reference genome. The resultant *juxtas*.*txt* file was used to create a graph of recombinant reads by placing the identified non-canonical read into a single 500-bp bin across the genome.

### Bacterial Strains and Culturing

Cyanobacterium *S. elongatus* UTEX 2973 was cultured under 100 or 200 µE white light at 38°C with 0.5% CO_2_ in 1X BG-11 medium (prepared from Sigma-Aldrich, Darmstadt, Germany, C3061-500ML). BW19851.*asd*::*apraR E. coli (10)* was cultured at 30°C in LB Lennox medium (Sigma, L3022-6X1KG). NEB 5-alpha *E. coli* (NEB, Ipswich, MA, C2987H), NEB 5-alpha F’I^q^ *E. coli* (NEB, Ipswich, MA, C2992I), and *P. putida* S12 were cultured in LB medium (Sigma-Aldrich, Darmstadt, Germany, L3522) at 37°C (*E. coli*) or 30°C (*P. putida*). When applicable, media were supplemented with 100 µg/mL diaminopimelic acid (DAP; Fisher AAB2239106) (*E. coli*), 50 µg/mL carbenicillin (Sigma C1389-5G; *E. coli*), 34 µg/ml of chloramphenicol (Teknova, Hollister, CA, C0322) (*E. coli*), 50 µg/mL kanamycin (American Bio, AB01100-00010; *S. elongatus*), or tetracycline (Teknova, Hollister, CA, T3325) at 10 µg/ml (*E. coli*) or 25 µg/ml (*P. putida*).

### Plasmid Construction

To generate plasmids for integrase testing in *P. putida* S12, DNA sequences consisting of a 600bp *attP* sequence (composed of the computationally determined *att* identity block flanked by GI DNA), a *tac* promoter, and the corresponding *int* were synthesized by Twist Biosciences. These sequences were then cloned into pACYC184 bearing additional I-SceI sites using NEBuilder HiFi Assembly mix (NEB, Ipswich, MA, E2621L). Integrase-testing plasmids were maintained with chloramphenicol selection in NEB 5-alpha F’I^q^ *E. coli*. Plasmids encoding frameshifted integrases were generated through either error-prone DNA synthesis or using inverse PCR to delete a nucleotide within the *int* sequence. Plasmids with mutant *attP* sequences contained 600 bp false *attP* sequences from *att*-flanking non-GI DNA.

To generate the positive control plasmid pUCP22-*tcR, tcR* (conferring tetracycline resistance) from pACYC184 was PCR amplified with Q5 DNA polymerase (NEB, Ipswich, MA, M0491S) and cloned into pUCP22 using restriction cloning (Eco53kI (NEB, Ipswich, MA, R0116S), XbaI (NEB, Ipswich, MA, R0145S), Monarch® Spin PCR & DNA Cleanup Kit (NEB, Ipswich, MA, T1130L), Ligase (NEB, Ipswich, MA,M0202L)). pUCP22-*tcR* was maintained in NEB 5-alpha *E. coli* with tetracycline selection. To verify all plasmid sequences, whole plasmid sequencing was performed by Plasmidsaurus (Louisville, KY) using Oxford Nanopore Technology with custom analysis and annotation.

Plasmids for integrase testing in *S. elongatus* UTEX 2973 were generated using Gibson assembly.

Constituent fragments were amplified from template plasmids: a backbone previously used for integrase-mediated cargo delivery in *S. elongatus* (pCargo (10)), the Sel_Y-Int_1 *int*, and an array of synthetic *attP* sites (Twist Biosciences). The assembly was performed with NEBuilder HiFi Assembly mix (NEB, Ipswich, MA, E2621L), and transformed into Lucigen EC100D pir+ cells (BioSearch Technologies, Petaluma, CA, ECP09500). Clones were sequence-verified via whole-plasmid sequencing by Quintara Biosciences.

### Integrase testing in *P. putida* S12 and KT2440

Electrocompetent *P. putida* S12 cells were prepared by washing subcultures (OD_600_ = 1.0) in 4°C 10% glycerol solution thrice with centrifugation steps occurring at 3000 x g for 15 minutes. All experiments were conducted from the same batch of electrocompetent cells stored at -80°C. 25 fmol of purified plasmid were electroporated (2500 V, 25 mF, 200 W, 2 mm gap cuvettes) into 50 ml of electrocompetent *P. putida* S12 or KT2440 in triplicate. Cells were recovered in LB broth for 2 hours at 30°C with aeration. Eight 10-fold serial dilutions of recovered cells were plated on LB with and without tetracycline. Plates were incubated for 18 hours and colony forming units (CFU) were measured. Percent Tc^R^ was calculated by dividing the number of tetracycline resistant CFU by the total CFU detected in the absence of Tc. Recombination efficiency was calculated by dividing the percent Tc^R^ following electroporation of integrase-testing plasmids by the percent Tc^R^ following electroporation of the positive control plasmid pUCP22-*tcR*.

To screen for correct insertion of plasmid sequences at the chromosomal *attB* site, colony PCR was conducted using OneTaq DNA polymerase (NEB, Ipswich, MA, M0486S) with one primer annealing to the plasmid DNA and its pair annealing to the *P. putida* genome adjacent to amplify across the *attR* site (**Supplemental Table 4**) (IDT, Coralville, IA). PCR products were run on an agarose gel for visualization with an Invitrogen 1 kb plus ladder (Invitrogen, Waltham, MA, 10787018) as a size standard.

### Serial Passaging

Tc^R^ colonies with integrated plasmids containing either Pal_S-Int_1 or Ppu_Y-Int_3 were cultured in LB medium without tetracycline in triplicate. Every 24 hours, cultures were diluted to an OD_600_ of 0.1 in 2 ml fresh LB medium. This passage was repeated thrice, allowing for 4 days of growth in medium without tetracycline selection. Cultures were then diluted and plated to achieve single colonies on LB agar without antibiotics. 72 colonies were then patched on LB agar containing tetracycline to identify the proportion of the population that had excised the integrated plasmid, and thus had lost Tc^R^.

To estimate the number of generations, the CFUs for cultures at an OD_600_ of 0.1 (0 h CFU) and cultures 24 hours later (24 h CFU) were calculated. The number of generations (n) occurring over 24 hours was calculated by n = (log(24 h CFU) - log(0 h CFU)) /log(2). This generation time was then multiplied by 4 to estimate the number of generations occurring over the 4 day period of serial passaging.

### Integrase testing in *S. elongatus*

Recipient *S. elongatus* was inoculated from a colony and grown to saturation in 25 mL BG-11 in a flask. Donors bearing the Sel_Y-Int_1 test plasmid, and a control strain bearing the same plasmid with a Bxb1 integrase, and the HB101 helper bearing pRL443 (29) were inoculated from glycerol stocks and grown overnight with carbenicillin supplementation. The next day, cultures were washed once in LB and resuspended at an OD_600_ of 10. Each donor was mixed in a 1:1 ratio with helper. Recipient was diluted to OD_750_ = 3 in nonselective BG-11 media, mixed in a 1:1 ratio with the donor-helper mixture, then spread on three autoclaved 25-mm HATF filters (Millipore, Darmstadt, Germany, HATF02500) on prewarmed BG-11 + 5% LB Lennox plates, for three replicates of each conjugation. Conjugations were incubated at 30°C in the dark at ambient CO_2_ overnight, then transferred to 38°C, 0.5% CO_2_, and 100 μE white light, for a total of 24 hours of conjugation. Filters were then transferred to 5 mL liquid BG-11 and vortexed to lift cells. Lifted exconjugants were concentrated via centrifugation, and spotted at various dilutions onto plates supplemented with kanamycin, wrapped in parafilm to retain moisture, and incubated at 38°C, 0.5% CO_2_, and 100 µE white light.

To test for integration of the Sel_Y-Int1 plasmid, kanamycin-resistant exconjugants were inoculated in 70 μL BG-11 media in a parafilm-wrapped 96-well V-bottom plate. Clones were grown for three days at 38°C, 0.5% CO_2_, and 200 μE white light, and then cultures were boiled at 65°C for 6 minutes followed by 95°C for 2 minutes. Boiled cells were subsequently screened for integration via PCR using Kapa 2G Fast master mix (Roche, Basel, Switzerland, KK5609) with primer pair OLQ1039/OLQ257 (targeting the locus with integrated plasmid) and OLQ1039/OLQ1038 (targeting the wild-type locus), cycling for 35 cycles with a 57°C annealing step and a 20s extension (**Supplemental Table 4**). Products were analyzed on a 1% agarose gel to determine presence or absence of integrated plasmid.

### Statistical Analyses

All integrase testing in *P. putida* and *S. elongatus* were conducted using three biological replicates. When present, error bars on graphs denote standard deviation.

## RESULTS

### Development of IOD software

The IOD software was developed to identify *attB* sites within bacterial genomes of interest to facilitate integrase-based integration of genetic cargo. It utilizes a list of integrase and *attB* sequence pairs derived from GIs predicted in our TIGER/Islander database of predicted GIs (20) (containing 182,734 *attBs* after deduplication based on sequence identity). The IOD software can act on a query genome sequence in two modes: *taxonomy* mode places the genome into the proper GTDB taxonomy (23) and identifies from close relatives integrase and *attB* pairs that are unoccupied by GIs in the query genome, as integrases from close relatives are more likely to function in the query genome. *Search* mode allows users to query user-provided (or all known) *attB* sites against their target genomes (**Figure 2**).

**Figure 2.**
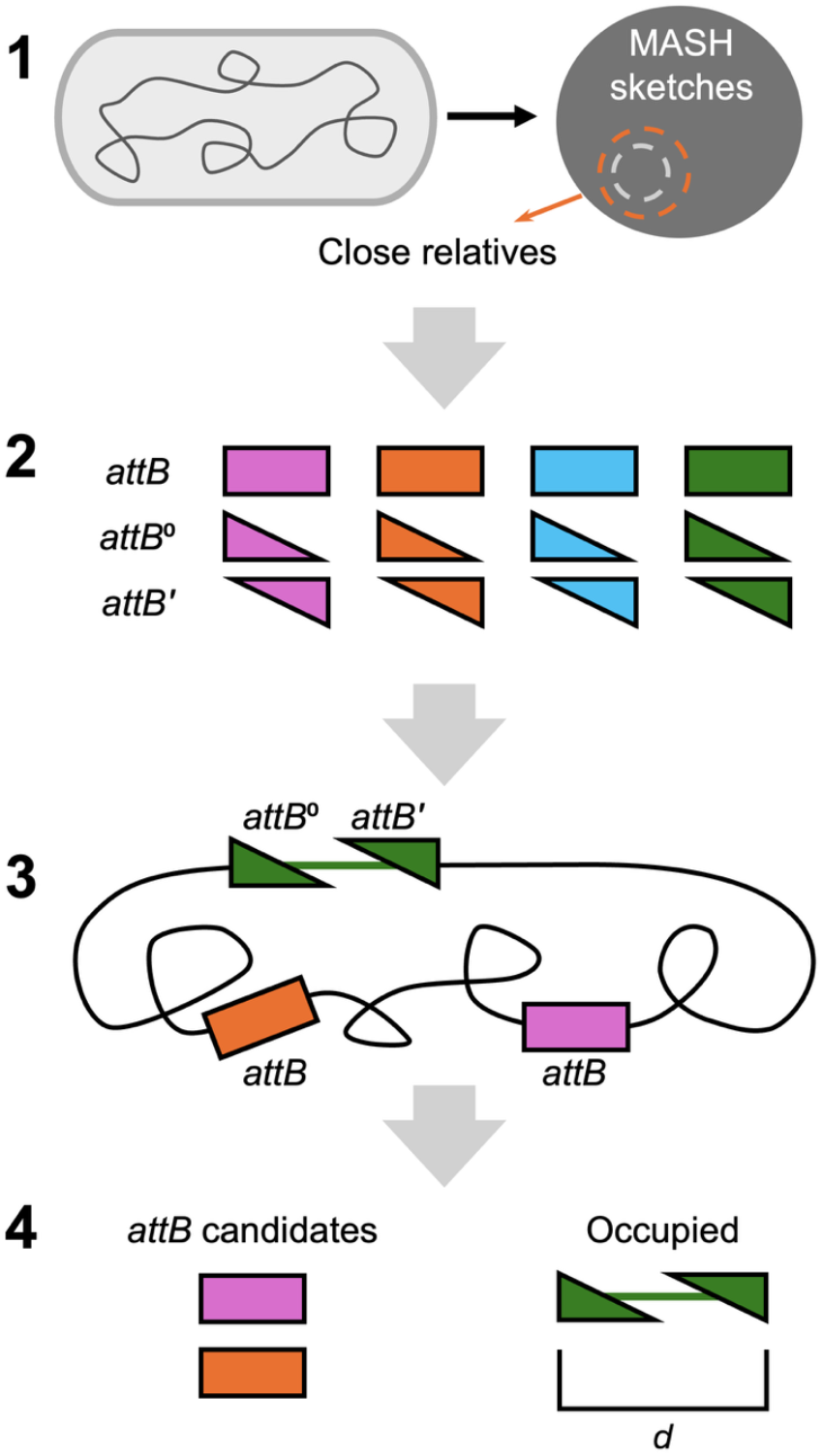
Workflow for IOD software in *taxonomic* mode. **1)** MASH (22) is used to determine close relatives of the input genome available in the IOD database. **2)** A query list of *attB*s and half attBs (see Methods; represented as triangles) are used as a query against the BLASTN database of the genome of interest. **3)** A series of filters locates candidate *attB* sites and collapses overlapping matches. **4)** GI-occupied *attBs* (identified by distance between hits to left and right *attB halves)* are rejected and the best candidate *attB* sites in the genome are reported separately from the duplicate sites.

To demonstrate broad potential utility of IOD for genome editing in phylogenetically diverse hosts, we have applied the IOD software to 183 genomes from a broad phylogenetic range of bacterial and archaeal species (**Table 1 and Supplementary Table 1**), including many relevant for biomanufacturing or medical applications (**Table 1**). Strains in **Table 1** typically contain numerous candidate *attB* sites identified using IOD in *taxonomy* mode, with only three exceptions (**Table 1**). However, most other species tested did not have candidate *attBs* predicted in *taxonomy* mode (97/142; **Supplementary Table 1**). For many phyla, this is caused by a lack of sequenced genomes from close relatives, resulting in too few *attB* search sequences. However, when running in *search* mode using all *attB* sequences in our database, only one strain out of the 183 genomes tested lacked an *attB* candidate; the total number of available *attB*s found in *taxonomic* mode was 234, rising to 1295 in *search* mode (**Supplementary Table 1**). There are no obvious trends as to which organisms are sources of the additional candidate *attB* sites found using *search* mode. In the cases we examined closely, source organisms span the entire phylogenetic tree with some crossing domains (e.g. integrase source is archaea and target was bacteria), further highlighting the utility of querying a large database for putative *attB* sites. These results indicate IOD software will find *attB*/integrase pairs for most bacteria and identify potential genomic locations for large DNA payload integration.

To benchmark software performance speed, we used the 183 genomes representing diverse bacterial and archaeal phyla (**Table 1 and Supplementary Table 1**). The average runtime per genome for both test-sets was 21.93 seconds, with no measurable impact of genome size (0.37-16Mbp) on runtime (**Supplemental Figure 2A**). Runtime is primarily dependent on the number of *attB* sequences used to query against the genome of interest (**Supplemental Figure 2B**). The maximum number of reference *attB* sequences collected from the closest relatives (based on phylogenomic comparison) of the input genome is 500 for *taxonomic* mode (run with default setting). However, there may be fewer *attB* sequences in the query depending on the species (see Methods), which results in a shorter runtime (e.g. *S. elongatus*). If the user runs the program in *search* mode without target sequences, the default database size is 182,734 (our entire *attB* database). The shortest allowable *attB* sequence in the database is 22 nucleotides (with default parameters). This was chosen to reduce false positives: given the probability of finding a random k-mer in a genome of length L is 2×L×4^-k^, the expected discovery rate for a random 22-mer in the *Pseudomonas putida* S12 genome is ∼3.67 × 10^-7^, and for at least one of 500 22-mers, 1.84 × 10^-4^. Benchmarking results emphasize the value of *taxonomic* mode for *attB* sequence identification given the low expected discovery rate and the correlation between runtime and number of query *attB* sequences. However, if computational resources or time are not of concern, *search* mode may be preferable.

### Prediction and Computational Analysis of IOD Integrases in *P. putida*

*P. putida* is a Gram-negative soil saprophytic bacterium that has been embraced as a biomanufacturing and synthetic biology chassis due to its hardiness, genetic tractability, and metabolic versatility (30). To refine our software predictions, we focused on the well-established laboratory strain *P. putida* S12. We predicted 37 candidate *attB* sites (**Supplementary Table 2**) and cross referenced with the TIGER/Islander predicted islands (**Supplementary Table 3**) to ensure that IOD differentiated *attB* sites (which would be available for integration of a synthetic cargo) from *attL/attR* sites (which already contain a genomic island and thus should not be provided to the user as available candidate *attB* sites). Unoccupied *attB*s are preferrable for biotechnology applications. Although, GI-occupied *attBs* can be utilized (and their repeated use is commonly observed in genomes, generating tandem GI arrays), the pre-existing integrases and/or its excisionases in a cell may destabilize a new insert that uses their *att* sites. Note that many GIs “repair” either the *attL* or *attR* to reconstitute the *attB*, a source of possible confusion (16,17).

**Table 2.**
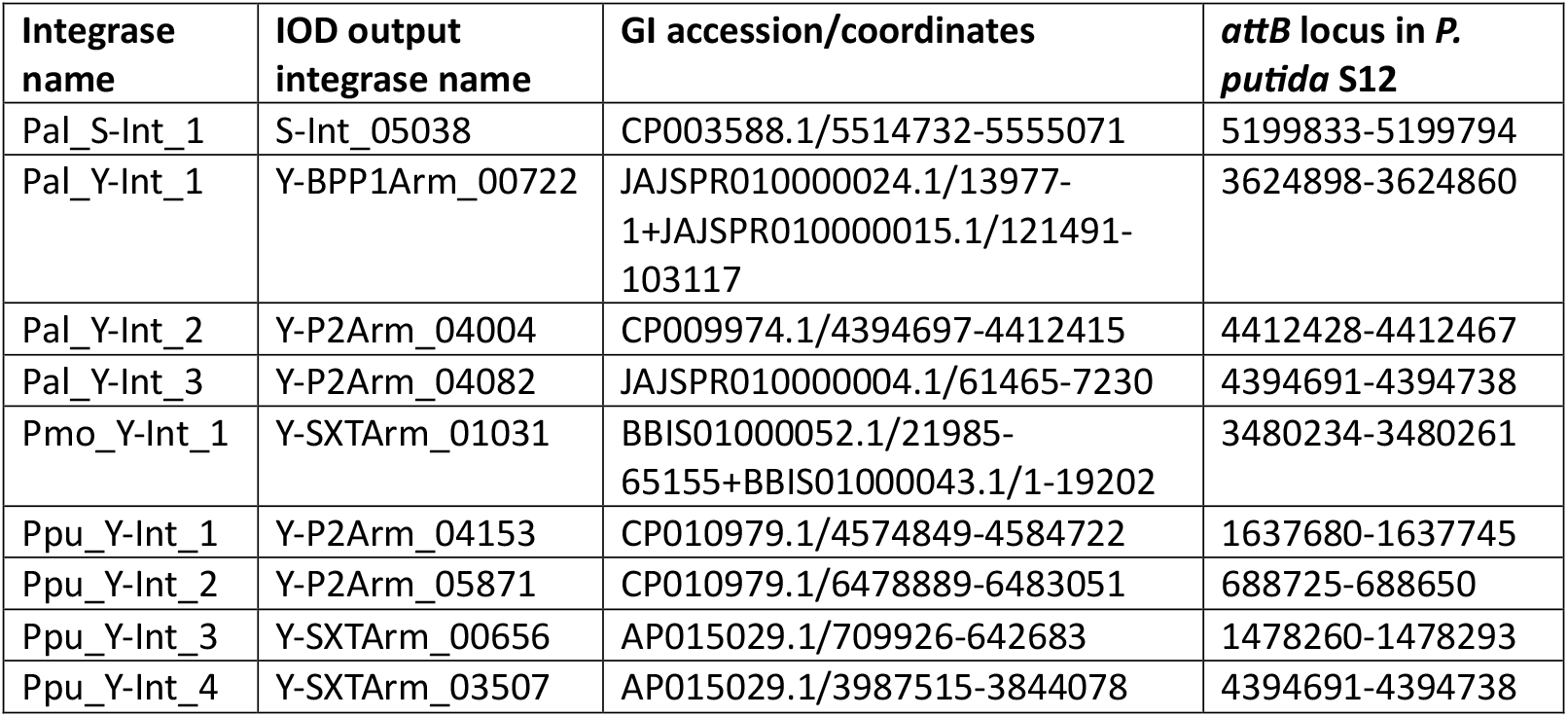
The nine tested integrase and *attB* pairs predicted for *P. putida* S12. Details on all IOD integrase candidates are shown in **Supplementary Table 2**.

The IOD software attempts to remove any spurious integrase/*attB* pairs as part of the upstream processing steps; however, additional verification may be necessary. In our initial test set for *P. putida* S12, we determined that the predicted Pmo_Y-Int_1 integrase was ∼125 aa shorter than expected (our predicted integrases had an average length of 393 aa, and the model tyrosine integrases are 356 aa [Lambda] and 343 aa [Cre]). We developed and implemented a filter to remove short integrases (see Methods), filtering out 315,597 of 923,744 unique tyrosine integrase sequences and 164,205 of 353,354 unique serine integrase sequences (mainly lacking the recombinase domain) from our initial integrase/*attB* site list.

To ensure we were testing functionally distinct integrases, we constructed two phylogenetic trees to determine the relationship between our IOD predictions for *P. putida* S12 (**Figure 3**). All predicted integrases were unique, except for three (Y-STXArm_01597, Y-STXArm_01983, Y-STXArm_02423). We chose to experimentally validate nine integrases from across the two trees (**Table 2, Figure 3**).

**Figure 3.**
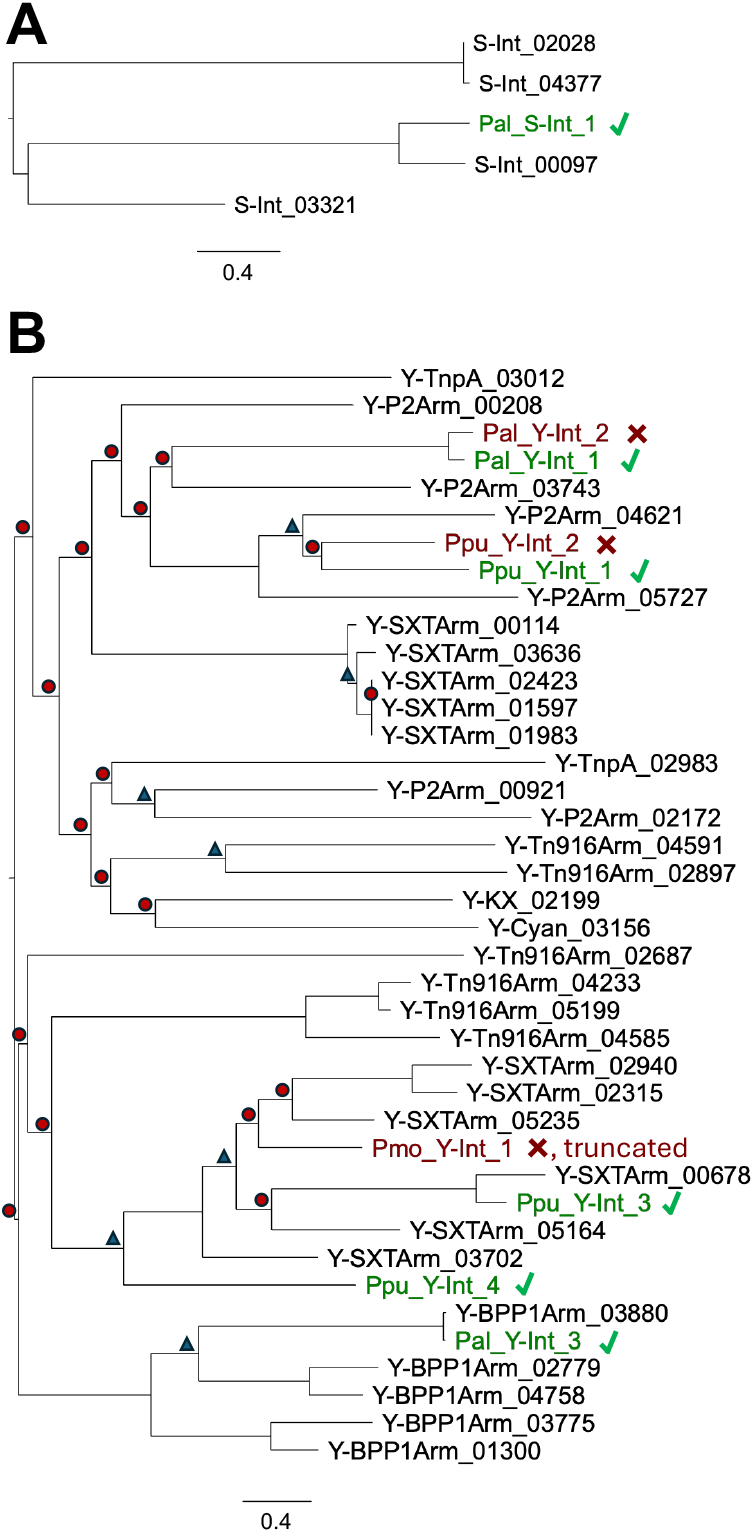
Phylogenetic tree of IOD predicted integrases for *P. putida* S12. Protein sequences were aligned to create a maximum likelihood tree for the **A)** Serine integrases (S-) and **B)** the Tyrosine integrases (Y-). Based on results from experimental validation, functional integrases are colored green and marked with a check and failed integrases are red colored with an X. Branch support values are ≥ 95% except for those marked with a triangle (50-94 %) or a circle (<50%).

We evaluated the integrase protein sequences to determine whether the catalytic serine or tyrosine residues were present. Previous work on phiC31 and Bxb1 serine integrases identified essential residues for enzymatic activity; these include a conserved N-terminal domain (31), the catalytic serine (S10 in Bxb1 (32) and S12 in phiC31 (33)), and conserved residues found in the active pocket required for catalytic activity (R8, Q22, D81, R82, L83, R85 in Bxb1; R10, Q26, R93, R96 in phiC31) (34). We confirmed that in all predicted serine integrases for *P. putida* S12, the catalytic serine residue was present, the N-terminal domain showed a high level of conservation, and the active pocket residues were fully conserved (**Supplemental Figure 3**).

In tyrosine integrases, the C-terminal catalytic domain is characterized by the crucial Tyr and the Arg-His-Arg catalytic triad (35). In all predicted tyrosine integrases, we identified the catalytic tyrosine (**Supplemental Figure 4**). We observed a few mutations in the catalytic triad in the integrases in this dataset. Two integrases had the first arginine residue of the catalytic triad mutated to a serine residue. In two integrases that were candidates for experimental validation (Pal_Y-Int_1 and Pal_Y-Int_2), the second arginine is mutated to a lysine. We also see a large clade of Y-BPP1 integrases (which includes Pal_Y-Int_3) missing the H308/H289 residue from the catalytic triad. We expect this to be a novel clade of integrases.

The lack of active recombination is an important criterion for selecting *attBs* for targeted integration because spurious excision or recombination activated by external stressors can reduce the stability of or damage the inserted genetic cargo. To test which predicted *attB* sites in *P. putida* S12 were optimal candidates for experimental validation of IOD, we treated *P. putida* S12 with MMC, a non-specific DNA damaging agent, which often induces GI excision or other recombination (potentially caused by transposons, homologous recombination, or non-homologous end joining (36,37)). Deep sequencing revealed that none of the *attB* sites that were selected for experimental validation in *P. putida* S12 were actively undergoing recombination (**Supplemental Figure 5**). While multiple regions in the *P. putida* S12 genome have sequencing reads indicating recombination above background levels, none of these correspond to predicted GIs (**Supplemental Table 3**), surprisingly, suggesting that the GIs in this strain are not excised upon MMC treatment.

### Experimental validation of predicted integrases for *Pseudomonas putida S12*

We constructed tetracycline resistance-conferring suicide vectors containing 600 bp *attP* sequences (**Supplementary Figure 6**) and the *tac* promoter driving expression of the cognate integrase (*int*) gene (**Figure 4A, Supplementary Table 5**). Because the vector does not replicate in *Pseudomonas*, plasmids that bear sequences encoding a nonfunctional integrase will be unable to recombine onto the chromosome and thus the plasmid, and the conferred tetracycline resistance, will be diluted out of the population. If the *int* sequence encodes a functional integrase and the predicted *attB* and *attP* are correct, the protein should mediate recombination between the plasmid *attP* sequence and the chromosomal *attB* sequence, stably conferring tetracycline resistance (Tc^R^) (**Figure 4B**).

**Figure 4.**
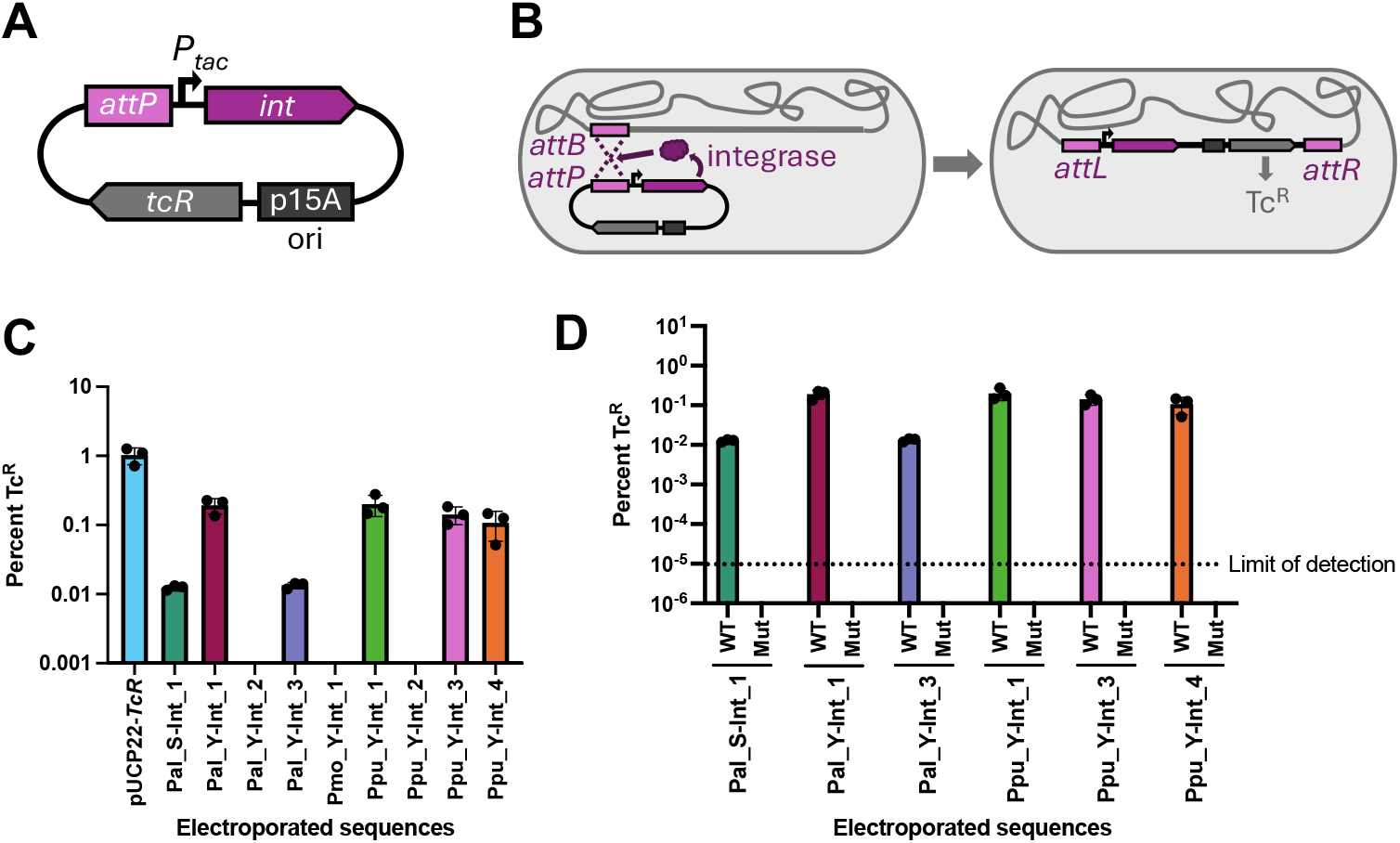
Using a suicide vector-based approach, 6/9 tested integrase and *attP* pairs mediate recombination in an integrase and *attP* sequence-specific manner. **A)** Diagram of an integrase testing plasmid. The testing plasmids bear the integrase gene (*int*) driven by a *P*_*tac*_ promoter, the 600 bp cognate *attP* sequence, a tetracycline resistance-conferring gene (*tcR*), and an origin that will not replicate in *Pseudomonas* (p15A ori). **B)** Schematic of integrase-mediated recombination between *attP* and *attB* sites that integrates the integrase testing plasmid sequences into the chromosome, conferring stable tetracycline resistance. **C)** The percent of the population with tetracycline resistance (Tc^R^) following electroporation of 25 fmol of an integrase test plasmid bearing the *int* and cognate *attP* of the listed integrase or a positive control, replicating plasmid, pUCP22-*TcR*. The limit of detection was 10^-5^ % TcR cells. **D)** Percent of the population with tetracycline resistance (Tc^R^) following electroporation of 25 fmol of (WT) a wild-type integrase test plasmid or (Mut) a plasmid containing either an incorrect *attP* sequence (Pal_Y-Int_1, Ppu_Y-Int_1, Ppu_Y-Int_3, Ppu_Y-Int_4) or a frameshifted *int* (Pal_S-Int_1, Pal_Y-Int_3).

Electroporation of plasmids bearing integrases Pal_S-Int_1, Pal_Y-Int_1, Pal_Y-Int_3, Ppu_Y-Int_1, Ppu_Y-Int_3, and Ppu_Y-Int_4 yielded an average of 0.01%, 0.19%, 0.01%, 0.20%, 0.14%, and 0.11%, Tc^R^ cells respectively (calculated by Tc^R^ CFU/total CFU)(**Figure 4C**). Therefore, these six integrases and *attP* pairs identified by the IOD software are capable of facilitating DNA recombination in *P. putida* S12. As expected, the heavily truncated Pmo_Y-Int_1 failed to yield Tc^R^ colonies, as did two additional integrases, Pal_Y-Int_2, and Ppu_Y-Int_2 integrase test plasmids into *P. putida* S12 yielded no Tc^R^ colonies, indicating that these three predicted integrases could not mediate recombination between the identified *attB* sites in our assay (**Figure 4C**). As expected, Pmo_Y-Int_1 failed likely due to the truncated protein produced.

Because acquiring stable Tc^R^ relies on successive successful electroporation followed by integrase-mediated recombination steps, to calculate integration efficiency we must account for electroporation efficiency. To assess the rate of plasmid uptake through electroporation for our population, we generated pUCP22-*tcR*, a tetracycline-resistance conferring, replicating plasmid that is similar in size to the *P. putida* S12 integrase-testing plasmids. Electroporation of 25 fmol of this plasmid yielded a population with 1.02% Tc^R^ cells, indicating that in our assay, electroporation will deliver a plasmid of nearly 6100 kb to 1/100 cells in the population. At this electroporation efficiency, integrases Pal_S-Int_1, Pal_Y-Int_1, Pal_Y-Int_3, Ppu_Y-Int_1, Ppu_Y-Int_3, and Ppu_Y-Int_4 facilitated recombination in an average of 1.2%, 18.7%, 1.3%, 19.4%, 13.8%, and 10.5% of the cells that had received the plasmid (**Supplementary Figure 7)**.

To verify that plasmid insertion into the chromosome is dependent on integrase-mediated recombination at *attB* sites and not mediated by other modes of recombination, we created additional plasmids with either 1) single nucleotide deletions in the *int* sequence to produce frameshifted, non-functional integrase proteins or 2) false *attP* sequences (see Methods). We observed that mutated versions of functional integrase/*attP* pairs did not generate Tc^R^ cells, and thus conclude that recombination is dependent on integrase activity and *attP* site sequence specificity (**Figure 4D**). To further verify that insertion is *attB* sequence-specific, we PCR amplified *attR* regions using a primer annealing to the plasmid backbone and a primer annealing to the chromosome (**Figure 5A**). For each integrase that mediated recombination, 8/8 screened Tc^R^ colonies yielded PCR products of the size expected for integration within the *attB* site, and no amplification was observed in the wild-type *P. putida* S12 negative control (**Figure 5B-G**). Therefore, the integrases are mediating recombination in an *attB* site-specific manner.

**Figure 5.**
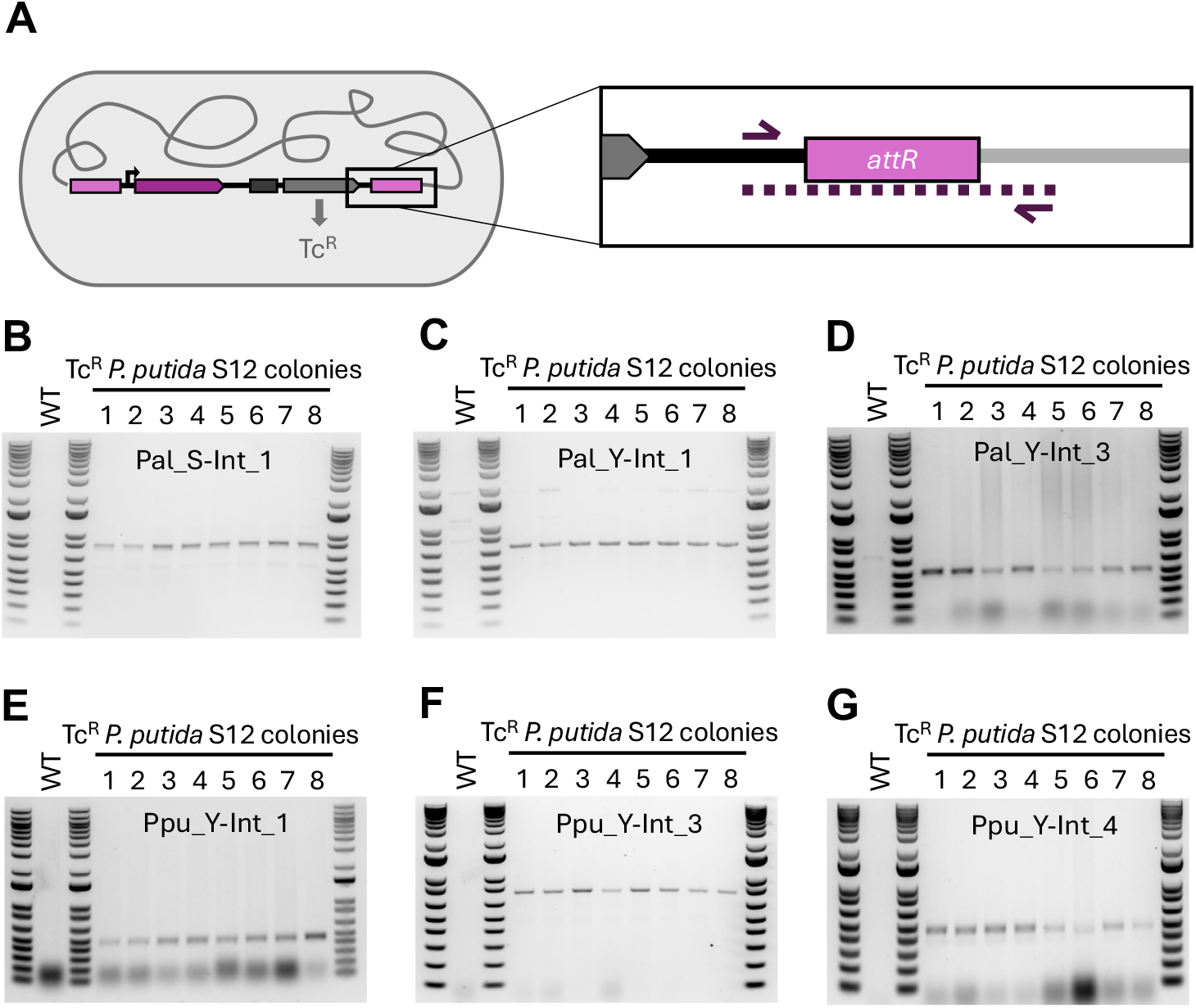
Functional integrases mediate site-specific recombination between predicted *attB* sites. **A)** To verify recombination occurs at the predicted chromosomal *attB* and plasmid *attP* sites, DNA from *attR*-containing junctions were PCR amplified using a primer annealing to chromosomal DNA and a primer annealing to plasmid DNA. **B-G)** PCR products using wild-type *P. putida* S12 (WT) or Tc^R^ *P. putida* S12 colonies post electroporation of *int* genes encoding **B)** Pal_S-Int_1, **C)** Pal_Y-Int_1, **D)** Pal_Y-Int_3, **E)** Ppu_Y-Int_1, **F)** Pal_S-Int_1, **G)** Ppu_Y-Int_4 and cognate *attP* containing test plasmids.

Integrase-inserted gene cargo should remain stably integrated when introduced unless cognate excisionases are present to allow the reverse reaction to occur, or there are other sources of genome instability. To test the stability of our integrated constructs, we serially passaged *P. putida* S12 with two integrated cargos (Pal_S-Int_1 and Ppu_Y-Int_3) in antibiotic-free medium every 24 hours for four days. By comparing growth on LB with and without tetracycline supplementation, we observed that the Pal_S-Int_1 test cargo remained stably integrated in the genome (0% Tc^S^), while the Ppu_Y-Int_3 test cargo excised in a small proportion of cells (1.39% Tc^S^) over approximately 26 generations. Excision may be due to induction of a cross-reactive excisionase (five putative excisionases are annotated in *P. putida* S12), or if the integrase is capable itself of catalyzing excision (i.e., bidirectional), or other processes such as endogenous recombination. Stability may be ensured, depending on causes of mobility by choosing the most stable integrase measured experimentally, or alternatively, maintaining (e.g., antibiotic) selection pressure. Cross-reacting excisionases can be addressed by changing the integrase choice or curing the strain of the offending interfering GI. Bidirectional integrases can also be identified experimentally using assays and excluded, or be tamed by providing *int* on a separate nonintegrating plasmid, or genetically inactivating the *int* once the construct is integrated.

We also used IOD to identify integrase and *attB* pairs for *P. putida* strain KT2440 (**Supplementary Table 2**), which is used more frequently than *P. putida* S12 in biomanufacturing applications (30). We observed that three of the six functional integrases tested in S12 were also predicted by the IOD software for KT2440 (Pal_Y-Int_1, Ppu_Y-Int_1, and Ppu_Y-Int_3; **Supplementary Table S2**). Using the same integrase test plasmids and assay, we demonstrated that all three integrases facilitated recombination in a sequence-specific manner, as evident from their generation of stably Tc^R^ colonies at 1.14%, 2.43%, and 2.39% of the total population respectively, and absence of colonies for the controls lacking integrase activity (**Supplemental Figure 8A and B**).

### Experimental validation of a predicted integrase in *S. elongatus*

To further demonstrate the potential utility of the IOD method, we sought to test a predicted integrase and *attP* pair for site-directed chromosomal integration in a cyanobacterial strain, *S. elongatus* UTEX 2973. Many cyanobacteria are notoriously difficult to engineer, due to their high copy number of chromosomes per cell (38,39), and slow chromosome segregation rates, so a tool for efficient genomic integration would be of great utility. We applied our IOD software to *S. elongatus* UTEX 2973 in *taxonomy* mode and discovered a single integrase and *attB* pair (**Supplementary Table 2**). To test this integrase in our target strain, we conjugated *S. elongatus* with a suicide vector bearing the lambda phage pR promoter-driven *int* encoding the Sel_Y-Int_1 integrase, *attP* site, and a kanamycin resistance marker to select for integrants (**Figure 6A and B**). As a negative control for the selection of colonies, we also conjugated, in parallel, the same construct with the Sel_Y-int_1 *int* replaced with the Bxb1 *int*, whose *attB* site is not found in the *S. elongatus* genome. Exconjugants from Sel_Y-int_1 grew on selective media, while the Bxb1 exconjugants did not (**Supplemental Figure 9**), indicating that survival of the Sel_Y-int_1 transformants was mediated by Sel_Y-int_1 activity towards its native *attB* site. To confirm that selected exconjugants had undergone integration of the test plasmid, we screened kanamycin-resistant colonies for integration at the native *attB* site via PCR. Because *S. elongatus* bears numerous chromosome copies, successful Sel_Y-Int_1-mediated recombination may occur at one or multiple *attB* loci in a single cell, resulting in a colony with a mixture of integrated and wild-type chromosome copies (“unsegregated”). All tested colonies demonstrated integration in some chromosome copies, and in most colonies, WT bands were very faint, suggesting almost complete segregation at the locus (**Figure 6C** and **Supplemental Figure 10**).

**Figure 6.**
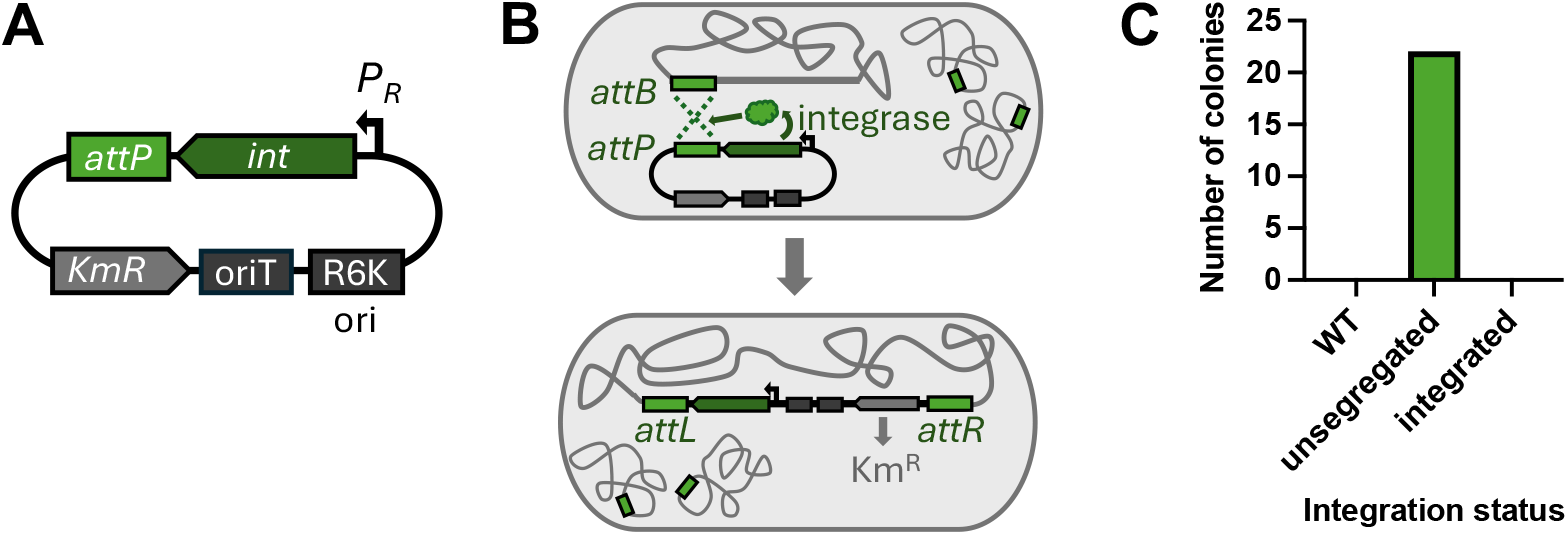
Testing Sel_Y-Int_1 integration at native cyanobacterial *S. elongatus attB* site. **A**) Schematic of the kanamycin resistance-conferring suicide plasmid bearing an R6K ori, origin of transfer (oriT) for conjugation, the Sel_Y-Int_1-encoding *int* driven by the lambda phage pR promoter, and the cognate *attP* site. **B**) Schematic of integrase-mediated recombination between the plasmid *attP* site and the many copies of the chromosomal *attB* site, which leads to stable kanamycin resistance. **C**) Km^R^ clones obtained were assayed for integration via PCR of the *attB* site. “Integrated” clones would have yielded a band only for the integrated plasmid, whereas “unsegregated” clones displayed both wild-type “WT” and integrated loci. WT clones would have displayed amplification for only the undisrupted locus, but no such instances were detected.

## DISCUSSION

Here, we have introduced IOD – a method for computationally identifying and experimentally utilizing native *attB* sites and their cognate integrases for genomic insertion of large DNA cargo. We have experimentally validated seven predictions in three biotechnologically relevant strains of bacteria (*P. putida* S12 and KT2440 and *S. elongatus* UTEX 2973), but IOD can be extended to a wide range of prokaryotes. These results illustrate the utility of our IOD method across phyla, and demonstrate use of both electroporation (*P. putida*) and conjugation methods (*S. elongatus*) for delivery of genetic cargo with these discovered integrases. Further, we tested integration of relatively small cargo sizes (∼6 kb in *P. putida* and 11 kb in *S. elongatus*); however, based on the size of naturally occurring GIs, we believe it likely that much larger cargos could generally be delivered using IOD-predicted integrases (6,40).

IOD also offers access of integrase-mediated genetic editing to a diversity of non-model microbes. When we applied our IOD software to multiple genomic sequences from 183 strains of bacteria and archaea spanning the tree of life, we predicted open *attB* sites for site-specific integrase-mediated cargo delivery with only one exception (182/183 tested genome sequences). The IOD software can search for candidate *attB* sites in any genomic sequence, including eukaryotes, though successes for non-prokaryotes may be infrequent due to low sequence similarity with the prokaryote-sourced *attB* sites. Further, utilizing integrase/*att* pairs that are drawn from genomes of near phylogenetic relatives to the strain of interest are less likely to require host factors or host background conditions not present in strain of interest, and may be more likely to work.

Integrases mediate stable DNA integration, but the plasticity of prokaryotic genomes may lead to loss of the inserted DNA cargo from the population over time. Our software aids in improving integration stability by filtering out *attB* sites present at multiple copies and those within GIs. To further ensure that IOD integration is site-specific and stable, we recommend that researchers do their due diligence, as done here by identifying recombination hotspots and, if possible, selecting *attB* sites that fall within stable regions of the genome. Additionally, promiscuous native integrases present in the target strain could, if actively produced, be cross reactive with the selected IOD *attP*. We suggest users compare amino acid sequences to ensure that the chosen IOD integrase is distinct from native integrases in the target strain. Finally, stability can be further improved by maintaining antibiotic selection to ensure that target cell populations maintain the inserted cargo. Future software improvements may include versions flagging integrases in site-promiscuous clades and those which likely disrupt essential genes (e.g., developmental genes (16)), to further improve output candidate integrases and *attB* pairs for genomic integration. However, systematic studies are required to establish reliable and efficient criteria for these searches.

By utilizing *attB* sites native to a host, IOD bypasses the requirement for multiple transformations or insertion of *attB* landing pads that are needed for traditional integrase-mediated genomic integration approaches (11-13). Establishing a new genetic engineering tool set in an undomesticated organism can take years of investment, but we were able to establish efficient IOD for three strains quickly. IOD’s simplicity requires knowledge of only: 1) a technique for introducing DNA into the target organism, 2) a selectable marker to screen for integration, and 3) a functional promoter for integrase expression, streamlining genome insertion in non-model organisms.

Serine integrases have traditionally been favored for biotechnological applications, due to their reduced requirements (e.g. short *att* sequences and no required cofactors) and integration stability compared to tyrosine integrases (5). Yet our test cases demonstrate that the tyrosine integrases can have great utility as well, mediating genomic integration at comparable rates to the tested serine integrase. Utilizing tyrosine integrases broadens the applicability of IOD, since tyrosine integrases are far more abundant in prokaryotic genomes than serine integrases (∼8-fold), greatly expanding the potential *attB* sites that may be present within a target organism. Furthermore, IOD enables selection of integrases from close relatives of a target strain, mitigating any potential dependencies that the integrases might have on host factors. Because of this, IOD extends the possibility of genetic engineering to virtually all prokaryotes to facilitate site-specific delivery of large cargo into the genome.

## DATA AVAILABILITY

All data underlying this article are available in the article or in the online supplementary material. All genomes processed are available at NCBI GenBank and the genome accession numbers are listed in Supplementary Table 1. The software is freely available at https://github.com/sandialabs/Integrase-On-Demand.

## ACKNOWLEDGEMENTS

We would like to thank Steve Branda and Brady Cress for thoughtful feedback on this manuscript, Grant Rybnicky (Northwestern University) for *P. putida* S12 and Anupama Sinha for assistance with sequencing. We also want to thank Jesse Cahill for his advice on integrase activity assays and help with construct design.

Sandia National Laboratories is a multi-mission laboratory managed and operated by National Technology & Engineering Solutions of Sandia, LLC (NTESS), a wholly owned subsidiary of Honeywell International Inc., for the U.S. Department of Energy’s National Nuclear Security Administration (DOE/NNSA) under contract DE-NA0003525. This written work is authored by an employee of NTESS. The employee, not NTESS, owns the right, title and interest in and to the written work and is responsible for its contents. Any subjective views or opinions that might be expressed in the written work do not necessarily represent the views of the U.S. Government. The publisher acknowledges that the U.S. Government retains a non-exclusive, paid-up, irrevocable, world-wide license to publish or reproduce the published form of this written work or allow others to do so, for U.S. Government purposes. The DOE will provide public access to results of federally sponsored research in accordance with the DOE Public Access Plan.

## AUTHOR CONTRIBUTIONS

Hannah M. McClain: Software, Formal Analysis, Funding acquisition, Visualization, Writing – original draft, Writing – review & editing. Lillian C. Lowrey: Methodology, Investigation, Visualization, Writing – original draft, Writing – review & editing. Laura B. Quinto: Methodology, Investigation, Visualization, Writing – original draft, Writing – review & editing. Ellis L. Torrance: Software, Formal Analysis, Writing – review & editing. Tomas R. Gagliano: Investigation, Writing – review & editing. Farren J. Isaacs: Funding acquisition, Supervision, Writing – review & editing. Joseph S. Schoeniger: Conceptualization, Funding acquisition, Project administration, Writing – review & editing. Kelly P. Williams: Conceptualization, Software, Data curation, Formal Analysis, Funding acquisition, Supervision, Writing – review & editing.

Catherine M. Mageeney: Conceptualization, Software, Data curation, Formal Analysis, Funding acquisition, Project administration, Visualization, Writing – original draft, Writing – review & editing.

## FUNDING

This work was supported by U. S. Department of Energy, Office of Science, through the Genomic Science Program, Office of Biological and Environmental Research, under the Secure Biosystems Design Science Focus Area, Intrinsic Control for Genome and Transcriptome Editing in Communities (InCoGenTEC) [24-026159 to J.S.S.]; Sandia National Laboratories Laboratory Directed Research and Development [222466 to K.P.W.]; U.S. Department of Energy, Office of Science, Office of Workforce Development for Teachers and Scientists (WDTS) under the Science Undergraduate Laboratory Internships Program (SULI) [to H.M.M]; and Phage Pathways: Reaching a New Energy Sciences Workforce (RENEW) supported by the Department of Energy Office of Science BER program [DE-SC0025673 to C.M.M]. Funding for open access charge: U. S. Department of Energy, Office of Science.

## CONFLICT OF INTEREST

No conflicts of interest.

## Notes

https://github.com/sandialabs/Integrase-On-Demand

### Competing Interest Statement

The authors have declared no competing interest.

