## Supplementary File for "Integrase-On-Demand: Bioprospecting integrases for targeted genomic insertion of genetic cargo"

### SUPPLEMENTARY MATERIALS

**Supplementary Table 1. IOD output for representatives from diverse GTDB phyla.**

The Diversity tab shows output information from both *taxonomic* mode and *search* mode for the 142 strains tested to span phylogenetic diversity. We list the GTDB domain, phylum, and species (names defined by GTDB v214 (23)), the 9-digit portion of the GenBank accession number from NCBI, genome size and number of contigs. The summarized output for the taxonomic and search modes include the number of *attB* sequence queries used, the number of candidate *attB* sites, which is further broken down into candidates in tRNA, occupied shows how many *attB* sites are currently occupied by GIs, runtime is the time it took IOD to run in seconds. The Health tab shows summarized output information for the 41 strains listed in **Table 1** from both *taxonomic* and *search* modes. This tab has the same structure as the Diversity tab without the GTDB defined domain and phylum.

**Supplementary Table 2. Candidate *attB* sites predicted by IOD**. *P. putida* S12 tab shows the candidate *attB* sites for *P. putida* S12. We also show two integrases that were experimentally tested but found on the occupied output list. *P. putida* KT2440 tab shows candidate *attB* sites for *P. putida* KT2440, candidates validated experimentally are bolded. *S. elongatus* tab shows the candidate *attB* sites from *S. elongatus* UTEX 2973. Each tab has the same organization: query contig is the contig from the target genome, strand dictates which strand the match is found on, query_coord is the coordinates of the *attB* sequence, query_attB is the sequence of the attB, integrase is the integrase name, islesID is the GI ID of the integrase/*att* pair source, ref_scaffold/coords is the GI accession and coordinates, source is which program called the GI, support is the support score for the GI, isles_type define which type of GI this was sourced from, ints,islesIDs are additional integrase island pairs that have the same *attB* sequence.

| Accession Number | Left Coord | Right Coord | Direction | GI Name | GI length | Type | Support Score |
|---|---|---|---|---|---|---|---|
| CP009974.1 | 1 | 2705 | - | 000495455.12.lexA\|IMS | 12011 | other | 33 |
| CP009974.1 | 1773566 | 1786082 | + | 000495455.13.T | 12517 | other | 5 |
| CP009974.1 | 2733967 | 2790931 | - | 000495455.57.nadE | 56965 | other | 83 |
| CP009974.1 | 2966334 | 2984279 | - | 000495455.18.Z\|C4 | 17946 | other | 49 |
| CP009974.1 | 4394697 | 4412415 | + | 000495455.18.P | 17719 | Phage2 | 9 |
| CP009975.1 | 1.1565 | 547421 | - | 000495455.42.hupB | 41702 | ICE1 | 4 |
| CP009975.1 | 13132 | 22537 | + | 000495455.9.IMS | 9406 | other | 4 |
| CP009975.1 | 53344 | 63613 | - | 000495455.10.hflC | 10270 | other | 1 |
| CP009975.1 | 268847 | 371193 | - | 000495455.102.HYP | 102347 | other | 2 |

**Supplementary Table 3. Genomic Islands predicted in S12.**

| Primer | Sequence | Purpose |
|---|---|---|
| OLQ1038 | ATCAACTTGCAAAAGGCACG | Assessing Sel_Y-Int_1 -mediated recombination in *S. elongatus* |
| OLQ1039 | GCGCTAGATCTTGCACATG | Assessing Sel_Y-Int_1 -mediated recombination in *S. elongatus* |
| OLQ257 | ACCATTGACATCACCATCCAG | Assessing Sel_Y-Int_1 -mediated recombination in *S. elongatus* |
| LL66 | GAATGGTCTTCGGTTTCC | Assessing int-mediated recombination in *P. putida* S12. Binds integrase test plasmid DNA to amplify *attR*. |
| LL73 | GATGATTGTTGAATTGGAGG | Assessing Pal_Y-Int_3-mediated recombination in *P. putida* S12. Binds *P. putida* chromosome to amplify *attR*. |
| LL74 | ATGAAGTCCCGAAAAACTC | Assessing Ppu_Y-Int_4-mediated recombination in *P. putida* S12. Binds *P. putida* chromosome to amplify *attR*. |
| LL115 | CTATTCCTCGCTCAGCT | Assessing Ppu_Y-Int_1-mediated recombination in *P. putida* S12. Binds *P. putida* chromosome to amplify *attR*. |
| LL113 | GTGAACAATCTCAGTCTTG | Assessing Ppu_Y-Int_3-mediated recombination in *P. putida* S12. Binds *P. putida* chromosome to amplify *attR*. |
| LL118 | TTGGCTGGGCGGGTT | Assessing Pal_Y-Int_1-mediated recombination in *P. putida* S12. Binds *P. putida* chromosome to amplify *attR*. |
| LL164 | CTTCGATGACGCAGACAA | Assessing Pal_S-Int_1-mediated recombination in *P. putida* S12. Binds *P. putida* chromosome to amplify *attR*. |

**Supplementary Table 4. Primers used in experiments**.

| Plasmid | Resistance | Backbone | Purpose |
|---|---|---|---|
| Int_att | Carb, Km | | Suicide vector for *attB* and integrase testing in *S. elongatus* |
| pLL06 | Tc, Cm | pACYC184 | Suicide vector for *attB* and Pal_Y-Int_3 integrase testing in *P. putida* S12 |
| pLL06mutint | Tc, Cm | pACYC184 | Suicide vector for *attB* and Pal_Y-Int_3.D169 integrase testing in *P. putida* S12 |
| pLL03 | Tc, Cm | pACYC184 | Suicide vector for *attB* and Pmo_Y-Int_1 integrase testing in *P. putida* S12 |
| pLL04 | Tc, Cm | pACYC184 | Suicide vector for *attB* and Ppu_Y-Int_2 integrase testing in *P. putida* S12 |
| pLL57 | Tc, Gentamycin | pUCP22 | Replicating vector to assess electroporation efficiency and Tc selection in *P. putida* S12 |
| pLL02 | Tc, Cm | pACYC184 | Suicide vector for *attB* and Ppu_Y-Int_4 integrase testing in *P. putida* S12 |
| pLL02mutatt | Tc, Cm | pACYC184 | Suicide vector for *attB* and Ppu_Y-Int_4.D214-215 integrase testing in *P. putida* S12 |
| pLL07 | Tc, Cm | pACYC184 | Suicide vector for *attB* and Ppu_Y-Int_1 integrase testing in *P. putida* S12 |
| pLL07mutatt | Tc, Cm | pACYC184 | Suicide vector for *attB* and Ppu_Y-Int_1 integrase testing in *P. putida* S12. Contains genomic, non-island DNA bounding that *attP* identity block. |
| pLL08 | Tc, Cm | pACYC184 | Suicide vector for *attB* and Ppu_Y-Int_3 integrase testing in *P. putida* S12 |
| pLL08mutatt | Tc, Cm | pACYC184 | Suicide vector for *attB* and Ppu_Y-Int_3 integrase testing in *P. putida* S12. Contains genomic, non-island DNA bounding that *attP* identity block. |
| pLL09 | Tc, Cm | pACYC184 | Suicide vector for *attB* and Pal_Y-Int_1 integrase testing in *P. putida* S12 |
| pLL09mutatt | Tc, Cm | pACYC184 | Suicide vector for *attB* and Pal_Y-Int_1 integrase testing in *P. putida* S12. Contains genomic, non-island DNA bounding that *attP* identity block. |
| pLL10 | Tc, Cm | pACYC184 | Suicide vector for *attB* and Pal_Y-Int_2 integrase testing in *P. putida* S12 |
| pLL11 | Tc, Cm | pACYC184 | Suicide vector for *attB* and Pal_S-Int_1 integrase testing in *P. putida* S12 |
| pLL11mutint | Tc, Cm | pACYC184 | Suicide vector for *attB* and Pal_S-Int_1.D931 integrase testing in *P. putida* S12. |

**Supplementary Table 5. Plasmids generated for integrase testing experiments.**

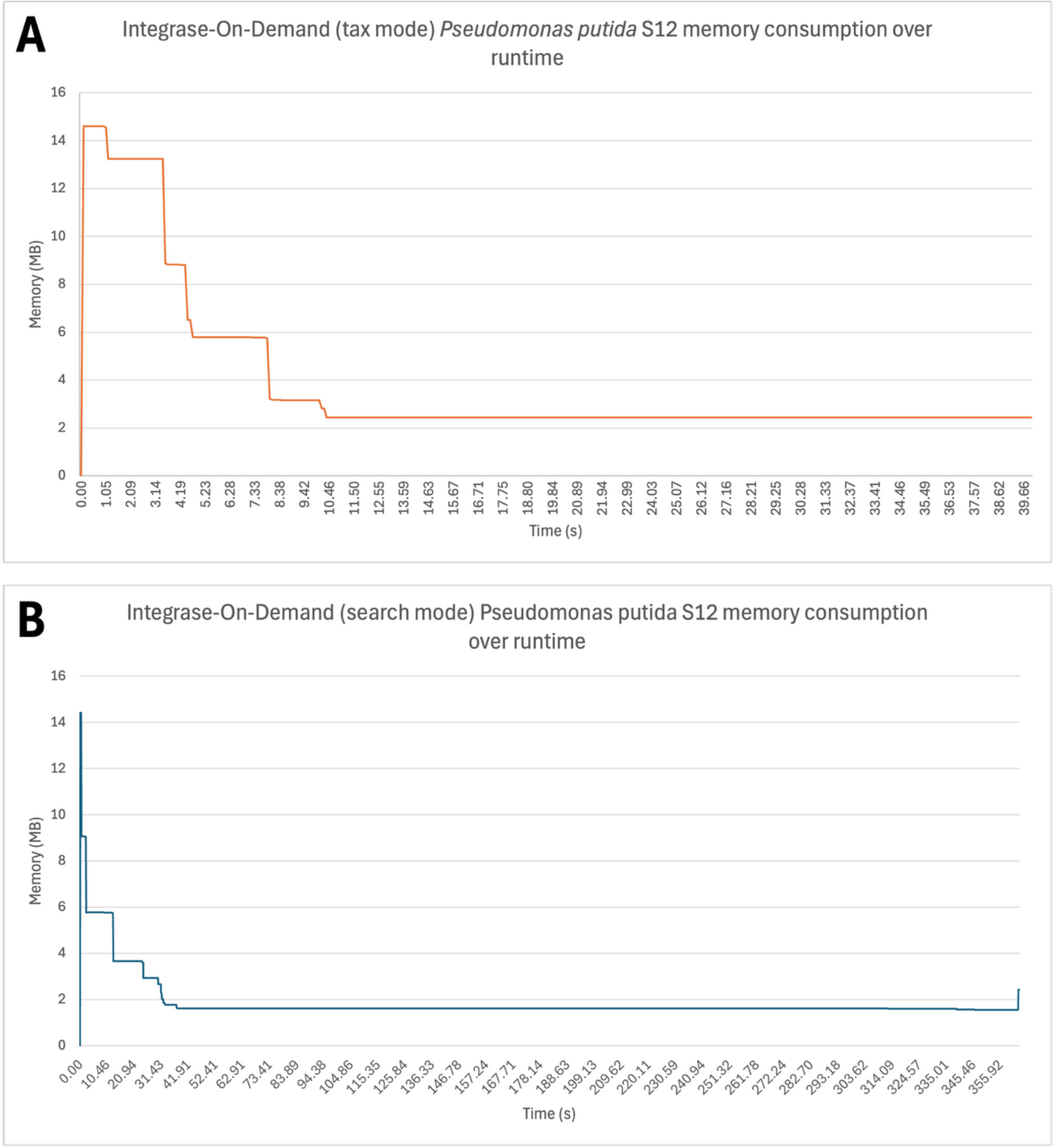


**Supplementary Figure 1. Comparing IOD memory consumption over runtime across two modes. A**) *P. putida* S12 genome in taxonomic mode using 500 *attB* sequences of close relatives as the query. **B)** *P. putida* S12 in search mode using all reference *attB* sequences as the query.

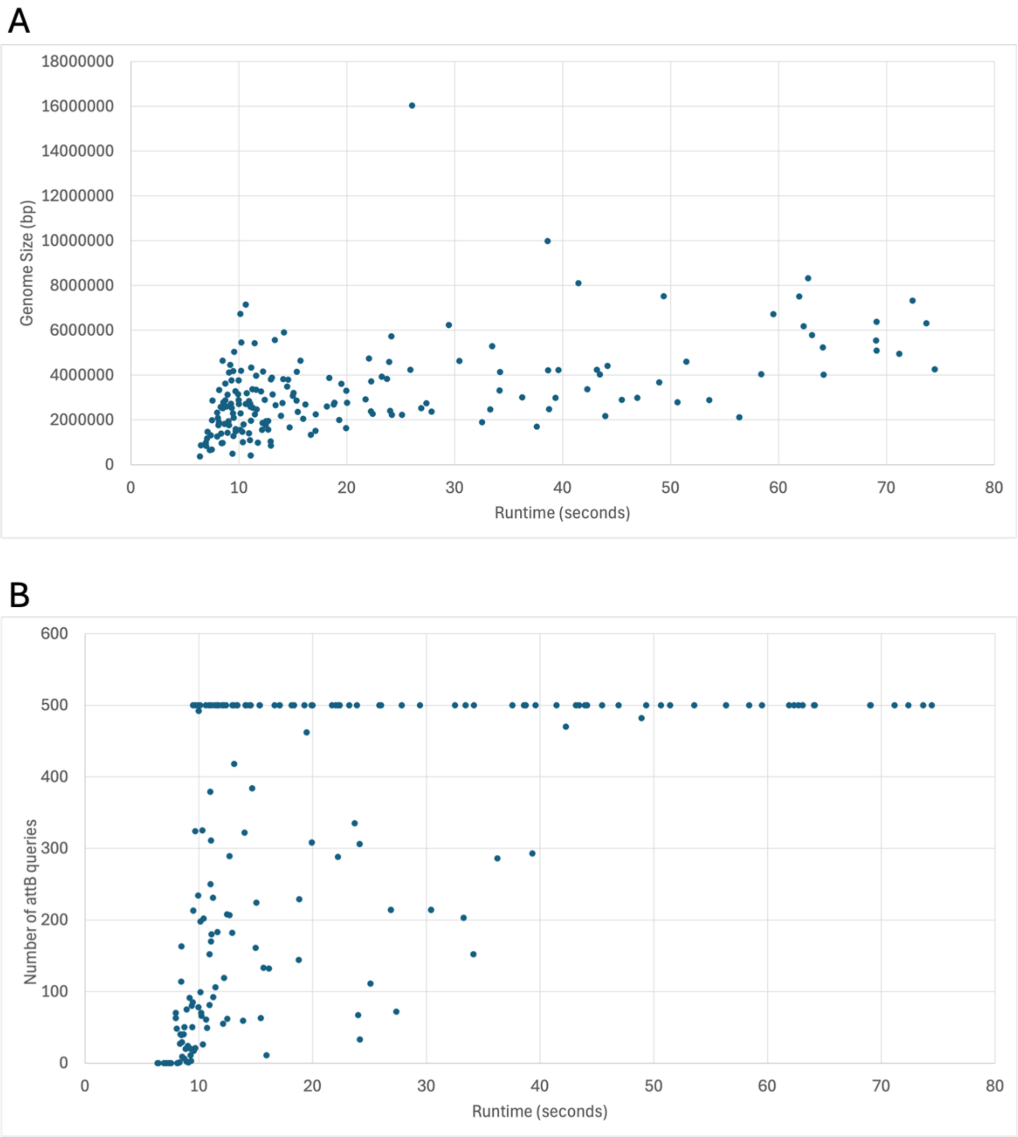


**Supplemental Figure 2. Run time is correlated with number of *attB* queries not genome size.** We applied IOD to 183 genomes across the phylogenetic tree of life and measured runtime. **A)** We examined the impact of genome size (y-axis) on the run time (x-axis). There is no correlation between genome size and run time. **B)** We investigated the number of *attB* queries (y-axis) versus run time (x-axis) and found a weak correlation between number of *attB* queries and run time.

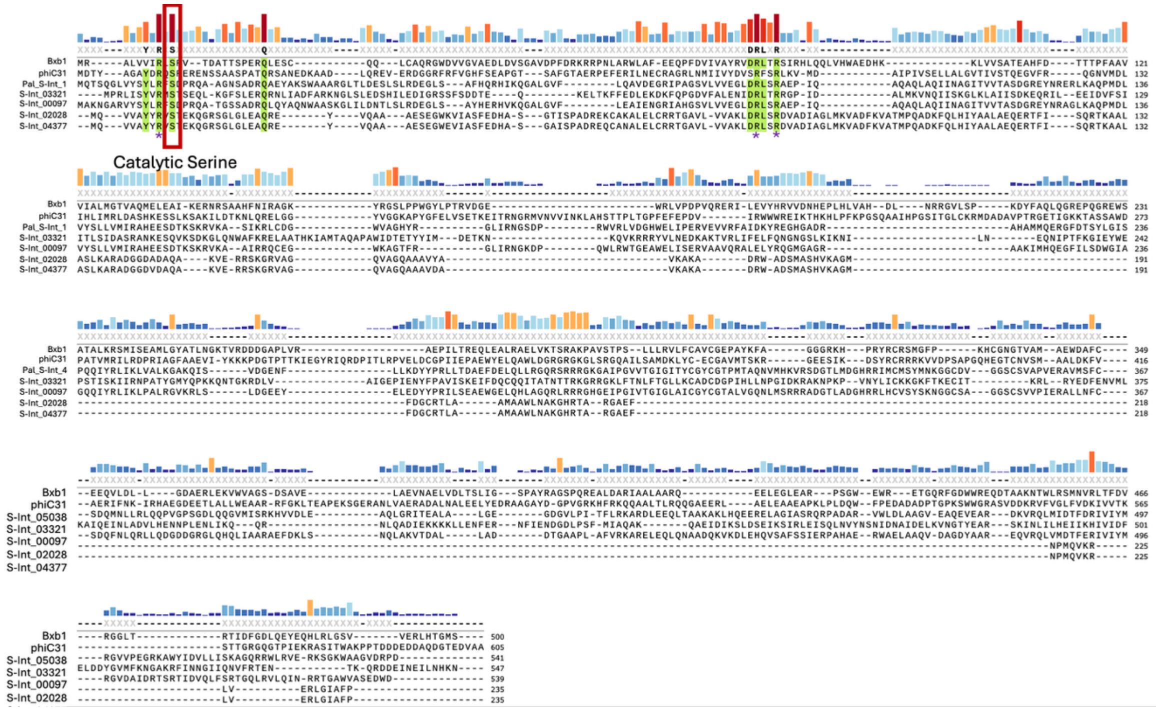

**Supplemental Figure 3. Serine Integrase Alignments.** The predicted serine integrases are listed with two model serine integrases, Bxb1 and phiC31. All serine integrases not experimentally tested are named as listed in **Supplemental Table 2**, experimentally verified serine integrases are names as in **Table 2**. The top bar graph shows the prevalence of each residue in the dataset. The catalytic serine residue is marked with a red box, and the other conserved residues are noted with a purple asterisk. Green highlights indicate residues found in >75% of the sequences.

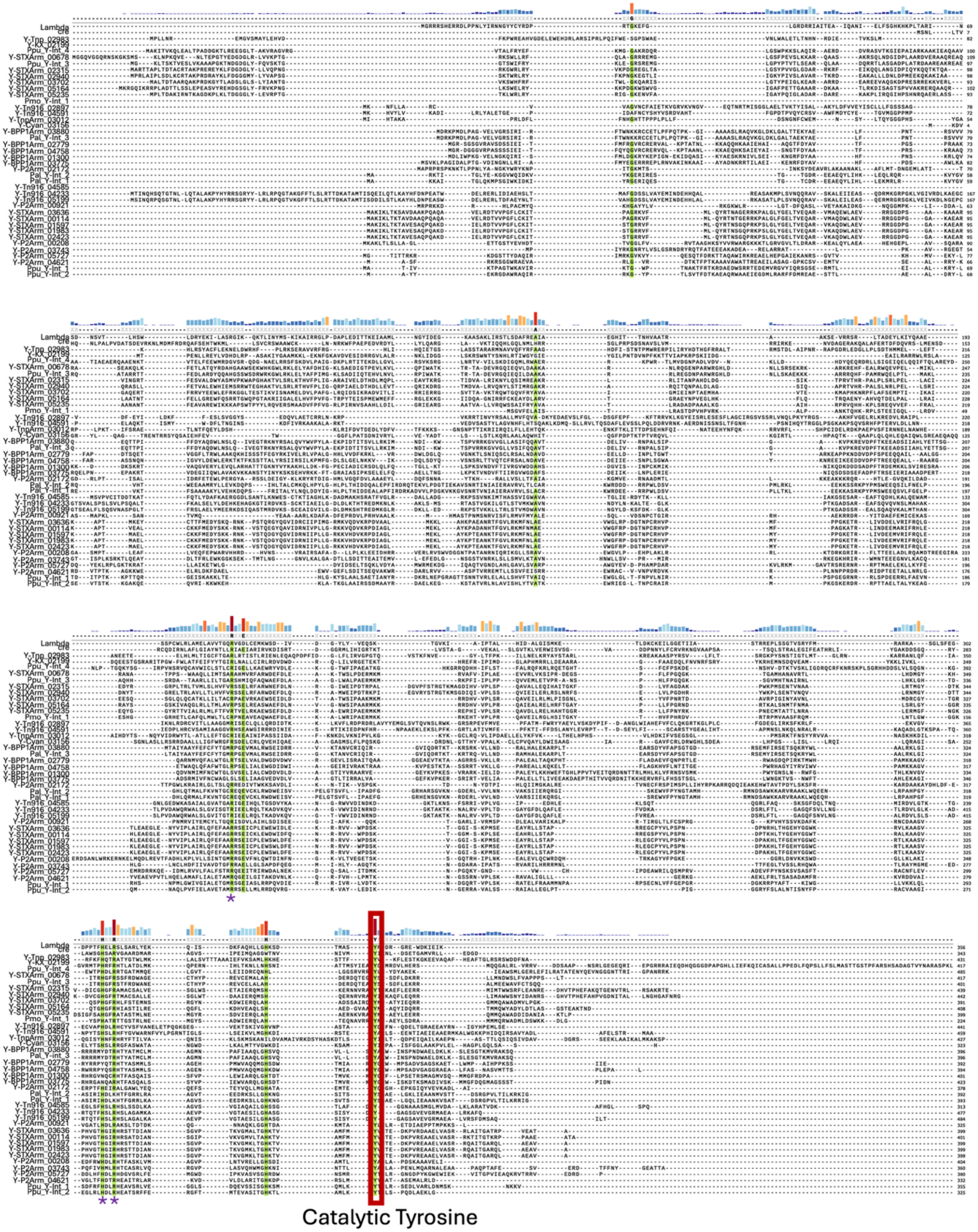
Catalytic Tyrosine

**Supplemental Figure 4. Tyrosine Integrase Alignments.** All predicted tyrosine integrases are listed with the model tyrosine integrases, Cre and Lambda. Names for tyrosine integrases experimentally verified are listed in **Table 2**, all non-experimentally verified tyrosine integrases are named as listed in **Supplemental Table 2**. The bar graph shows the prevalence of each amino acid at that location. The catalytic tyrosine is highlighted in the red box. The catalytic triad residues are noted with a purple asterisk. Any residue found in >75% of the sequences is highlighted green.

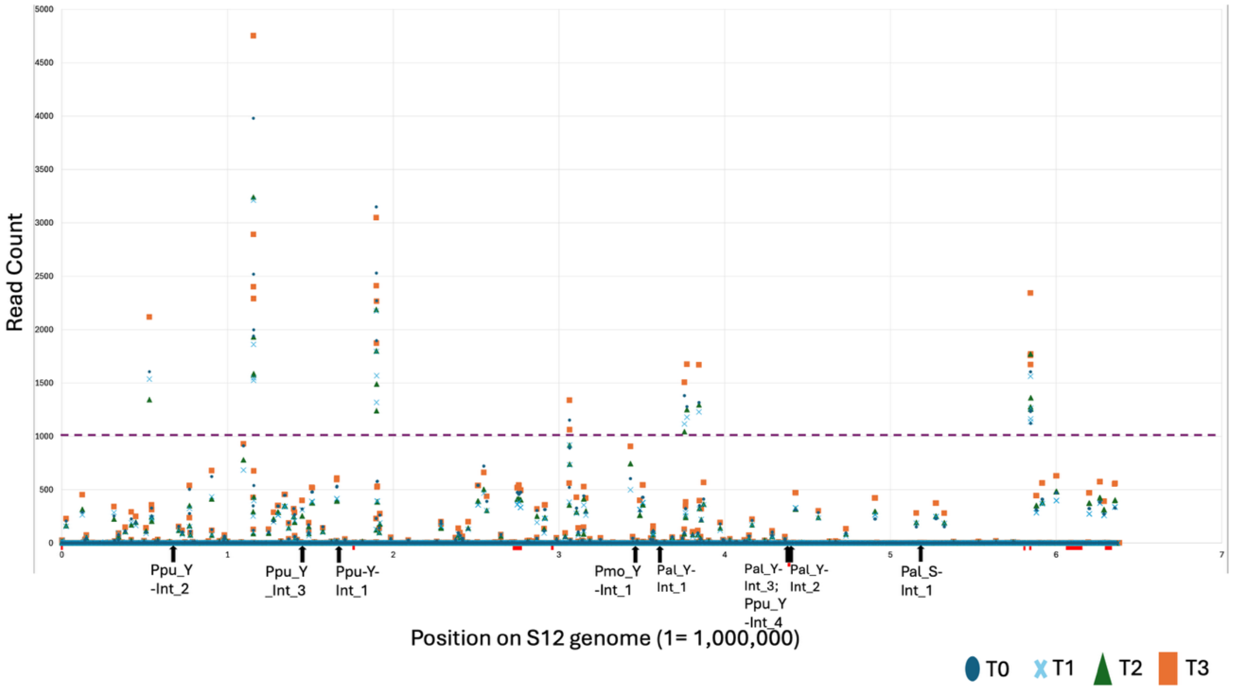


**Supplemental Figure 5. Graph of all recombinant reads of *P. putida* S12**. The number of recombinant reads (y-axis) in each 500bp bin after deep sequencing and analysis with Juxtaposer are noted for each time point post MMC addition. The x-axis is the position along the genome in *P. putida* S12. The integrase *attB* sites are noted with black arrows. GIs are marked with red boxes along the X-axis. Anything below the threshold of 1000 reads is within the noise of the dataset.

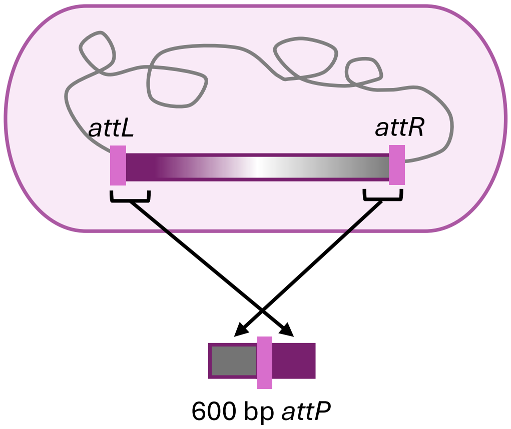


**Supplemental Figure 6. Schematic of *attP* sequence determination**. 600 bp *attP* sequences were built for each integrase test plasmid by bounding the *att* identity block with flanking island DNA to reconstruct the original island *attP.*

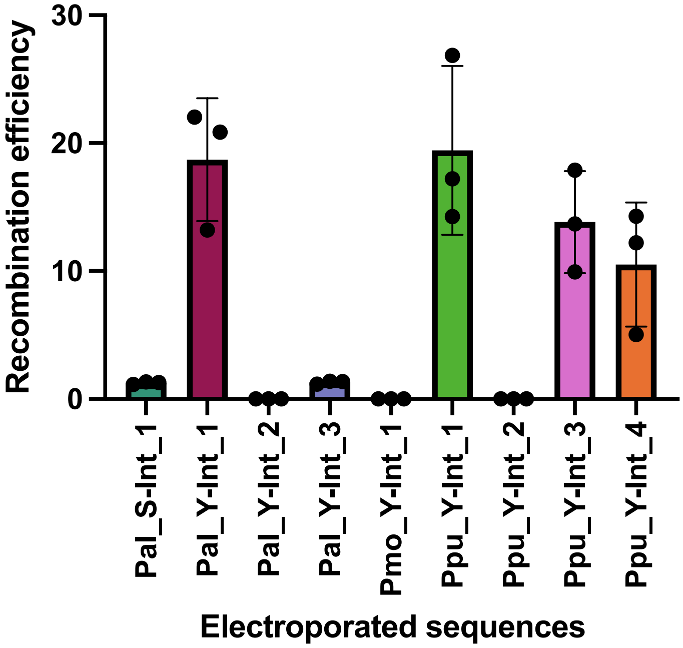


**Supplementary Figure 7. Integrases efficiently mediate recombination introducing genetic cargo into the *P. putida* S12 genome.** Efficiency was calculated by dividing the percentage of $Tc^R$ cells in the population following electroporation of each integrase test plasmid by the percentage of $Tc^R$ cells following electroporation of the replicating, positive control plasmid pUCP22-*tcR*

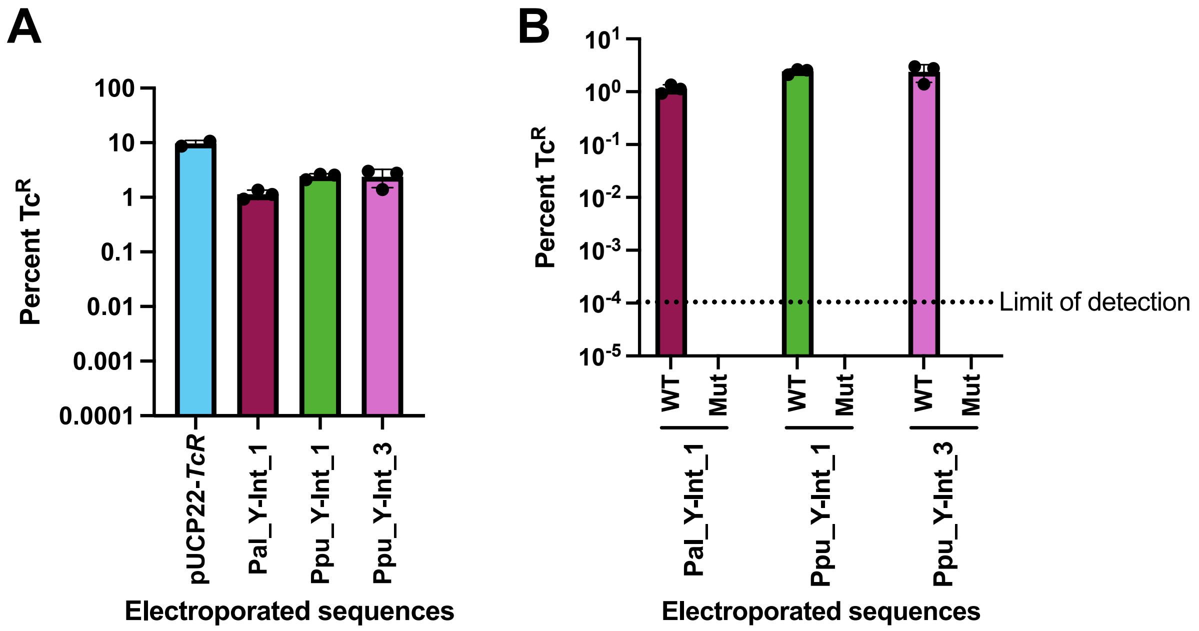


**Supplemental Figure 8. Three integrases predicted by IOD function in *P. putida* KT2440. A)** The percent of the population with tetracycline resistance ($Tc^R$) following electroporation of 25 fmol of an integrase test plasmid bearing the *int* and cognate *attP* of the listed integrases (Pal_Y-Int_1, Ppu_Y-Int_1, Ppu_Y-Int_3) or a positive control, replicating plasmid, pUCP22-*TcR*. **B)** The percent of the population with tetracycline resistance ($Tc^R$) following electroporation of 25 fmol of (WT) a wild-type integrase test plasmid or (Mut) a plasmid containing a false *attP* sequence. The minimum limit of detection for $Tc^R$ cells was $10^{-4}$ percent.

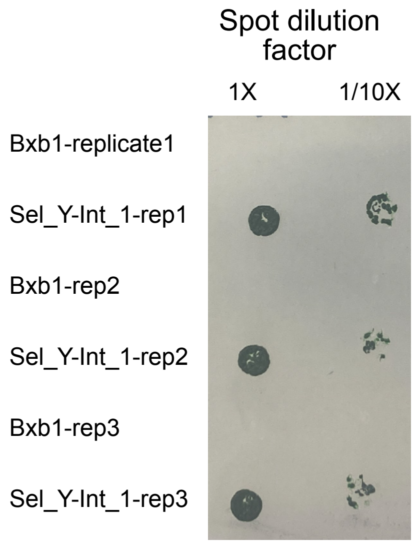


**Supplemental Figure 9. Integrase-mediated genomic insertion in *S. elongatus* requires an *attB* site**. Cells from each conjugation with the experimental Sel_Y-int_1 integrase, whose *attB* site is natively present in the *S. elongatus* genome, as well as a negative control with the Bxb1 integrase, were spotted on selective media. Each replicate is the result of a separate conjugation from the same initial recipient, helper, and donor cultures (see Methods).

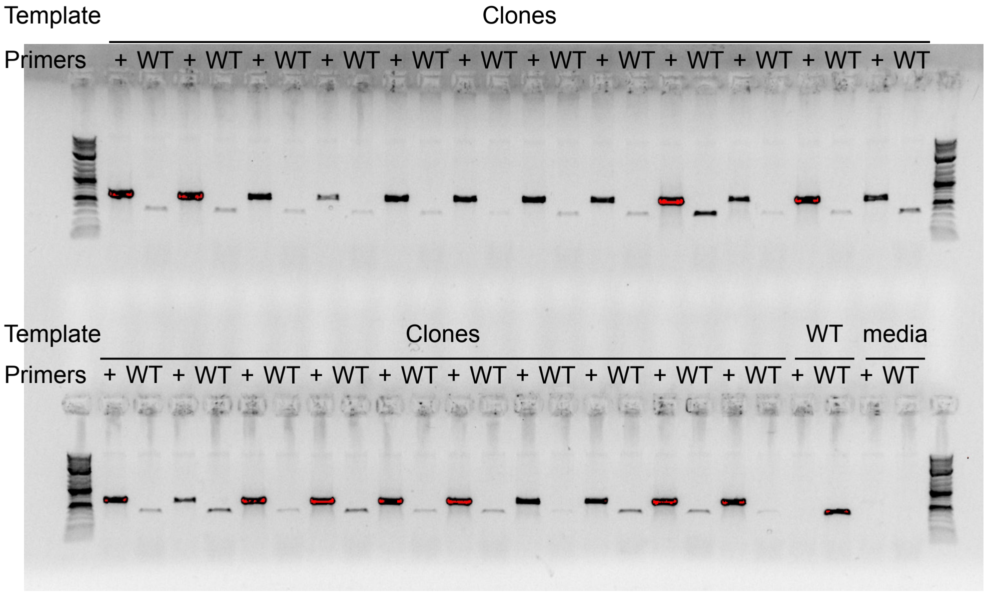


**Supplemental Figure 10. PCR verification of Sel_Y-int_1-mediated genomic insertion**. Colonies obtained from conjugation with Sel_Y-int_1 were inoculated and assayed via PCR for integration. Primers in the "+" PCR condition detect integration of the plasmid at the predicted *attB* site, and the "WT" PCR primers detect the undisrupted *attB* site. Each pair of "+" and "WT" lanes correspond to a single clone. Controls were performed with wild-type cells ("WT") and media-only template.
