## Supplementary Table 1 for "Integrase-On-Demand: Bioprospecting integrases for targeted genomic insertion of genetic cargo"

| | | | | | | *Taxonomic Mode* | | | | | *Search Mode* | | | | |
|---|---|---|---|---|---|---|---|---|---|---|---|---|---|---|---|
| **GTDB domain** | **GTDB phylum** | **GTDB species** | **GCA** | **Genome size (bp)** | **Contigs** | **No. attB queries** | **attB candidates** | **attBs in tRNA genes** | **Occupied** | **Runtime (s)** | **No. attB queries** | **attB candidates** | **attBs in tRNA genes** | **Occupied** | **Runtime (s)** |
| Archaea | Aenigmatarchaeot | JAGLWW01__sp020355185 | 020355185 | 843290 | 1 | 0 | 0 | 0 | 0 | 6.95 | 182734 | 1 | 0 | 4 | 82.63 |
| Archaea | Altiarchaeota | IMC4__sp001742785 | 001742785 | 1390916 | 1 | 27 | 0 | 0 | 0 | 8.33 | 182734 | 0 | 0 | 8 | 87.91 |
| Archaea | Asgardarchaeota | PR6__sp021513715 | 021513715 | 3368368 | 1 | 231 | 0 | 0 | 1 | 11.25 | 182734 | 1 | 0 | 32 | 122.74 |
| Archaea | B1Sed10-29 | WALL01__sp023129115 | 023129115 | 1263008 | 19 | 70 | 0 | 0 | 0 | 7.97 | 182734 | 1 | 0 | 17 | 105.03 |
| Archaea | EX4484-52 | JAHYJR01__sp019429005 | 019429005 | 1092481 | 15 | 379 | 0 | 0 | 3 | 11.01 | 182734 | 3 | 1 | 45 | 118.14 |
| Archaea | Hadarchaeota | DG-33__sp004375695 | 009619155 | 1310696 | 20 | 0 | 0 | 0 | 0 | 7.38 | 182734 | 2 | 0 | 23 | 100.42 |
| Archaea | Halobacteriota | Halohasta__litchfieldiae | 002788215 | 3373673 | 1 | 470 | 4 | 2 | 2 | 42.27 | 182734 | 4 | 1 | 20 | 108.31 |
| Archaea | Huberarchaeota | Huberarchaeum__crystalense | 001872885 | 416970 | 26 | 311 | 0 | 0 | 0 | 11.07 | 182734 | 2 | 1 | 40 | 77.98 |
| Archaea | Hydrothermarchae | Hydrothermarchaeum__profundi | 002011125 | 2087920 | 22 | 48 | 0 | 0 | 0 | 8.05 | 182734 | 1 | 1 | 46 | 117.18 |
| Archaea | Iainarchaeota | Forterrea__multitransposorum | 009392915 | 850128 | 1 | 500 | 0 | 0 | 2 | 12.95 | 182734 | 1 | 0 | 16 | 115.8 |
| Archaea | Methanobacteriot | Methanococcus__maripaludis | 003052125 | 1759327 | 1 | 208 | 2 | 2 | 1 | 12.48 | 182734 | 3 | 2 | 27 | 122.26 |
| Archaea | Methanobacteriot | Thermococcus__gammatolerans | 000022365 | 2045439 | 1 | 11 | 1 | 1 | 0 | 15.95 | 182734 | 2 | 1 | 6 | 97.58 |
| Archaea | Micrarchaeota | Mancarchaeum__acidiphilum | 002214165 | 964161 | 1 | 40 | 0 | 0 | 0 | 8.39 | 182734 | 1 | 0 | 6 | 79.04 |
| Archaea | Nanoarchaeota | GCA-020343435__sp020343435 | 020343435 | 655986 | 1 | 0 | 0 | 0 | 0 | 7.28 | 182734 | 1 | 1 | 6 | 80.45 |
| Archaea | Nanohaloarchaeot | SW-4-43-9__sp003009795 | 003009795 | 968507 | 1 | 163 | 0 | 0 | 0 | 8.47 | 182734 | 2 | 0 | 4 | 91.76 |
| Archaea | SpSt-1190 | JAAOXX01__sp018220935 | 018220935 | 861093 | 37 | 0 | 0 | 0 | 0 | 6.46 | 182734 | 2 | 1 | 43 | 89.05 |
| Archaea | Thermoplasmatota | Thermoplasma__acidophilum | 000195915 | 1564907 | 1 | 289 | 0 | 0 | 0 | 12.7 | 182734 | 3 | 1 | 8 | 82.71 |
| Archaea | Thermoproteota | Nitrosocosmicus__franklandus | 900696045 | 2871903 | 1 | 6 | 0 | 0 | 0 | 8.73 | 182734 | 2 | 0 | 29 | 111.25 |
| Archaea | Undinarchaeota | Undinarchaeum__sp002687935 | 002687935 | 674879 | 11 | 0 | 0 | 0 | 0 | 7.48 | 182734 | 4 | 3 | 5 | 75.7 |
| Bacteria | 4484-113 | JAHEFU01__sp022646745 | 022646745 | 4163665 | 1 | 119 | 0 | 0 | 1 | 12.21 | 182734 | 14 | 9 | 25 | 130.55 |
| Bacteria | AABM5-125-24 | JABWCQ01__sp020637445 | 020637445 | 3130534 | 4 | 75 | 1 | 1 | 1 | 8.93 | 182734 | 5 | 3 | 18 | 118.69 |
| Bacteria | Acidobacteriota | JAAYLR01__sp016699405 | 016699405 | 5289494 | 1 | 500 | 1 | 0 | 2 | 33.44 | 182734 | 12 | 4 | 37 | 217.34 |
| Bacteria | Actinomycetota | Streptomyces__yunnanensis | 016917755 | 9980755 | 1 | 500 | 14 | 8 | 6 | 38.58 | 182734 | 42 | 17 | 64 | 620.82 |
| Bacteria | Aerophobota | JAGHCO01__sp021160405__X1 | 000404365 | 368051 | 32 | 0 | 0 | 0 | 0 | 6.37 | 182734 | 5 | 3 | 6 | 80.03 |
| Bacteria | Aquificota | Hydrogenobaculum__sp000213785 | 000213785 | 1552776 | 1 | 55 | 0 | 0 | 0 | 12.12 | 182734 | 7 | 3 | 18 | 111.25 |
| Bacteria | Armatimonadota | JALHNM01__sp019454125 | 019454125 | 2685503 | 1 | 144 | 2 | 2 | 0 | 18.78 | 182734 | 15 | 4 | 16 | 127.76 |
| Bacteria | Atribacterota | Atribacter__laminatus | 015775515 | 3139991 | 1 | 418 | 1 | 0 | 0 | 13.1 | 182734 | 8 | 6 | 21 | 131.15 |
| Bacteria | Auribacterota | JACKPL01__sp024233545 | 024233545 | 2471892 | 26 | 183 | 0 | 0 | 0 | 11.62 | 182734 | 7 | 4 | 45 | 124.97 |
| Bacteria | Bacillota | Streptococcus__pneumoniae | 001457635 | 2110969 | 1 | 500 | 44 | 3 | 16 | 56.34 | 182734 | 95 | 3 | 20 | 282.88 |
| Bacteria | Bacillota_A | Caproicibacterium__sp902809935 | 902809935 | 2365056 | 1 | 500 | 0 | 0 | 0 | 27.84 | 182734 | 11 | 4 | 14 | 149.34 |
| Bacteria | Bacillota_B | Desulfitobacterium__hafniense | 000010045 | 5727535 | 1 | 306 | 8 | 4 | 1 | 24.12 | 182734 | 14 | 7 | 19 | 129.53 |
| Bacteria | Bacillota_C | Acidaminococcus__intestini | 000411395 | 2287857 | 1 | 99 | 6 | 1 | 2 | 10.14 | 182734 | 21 | 9 | 16 | 149.12 |
| Bacteria | Bacillota_D | Natranaerofaba__carboxydovora | 022539405 | 3305542 | 1 | 308 | 0 | 0 | 0 | 19.93 | 182734 | 5 | 3 | 27 | 127.48 |
| Bacteria | Bacillota_E | R501__sp902809825 | 902809825 | 2602467 | 1 | 500 | 2 | 1 | 0 | 18.14 | 182734 | 11 | 5 | 17 | 183.47 |
| Bacteria | Bacillota_F | Halothermothrix__orenii | 000020485 | 2578147 | 1 | 1 | 0 | 0 | 1 | 8.29 | 182734 | 8 | 4 | 28 | 129.86 |
| Bacteria | Bacillota_G | Limnochorda__pilosa | 001544015 | 3817037 | 1 | 500 | 0 | 0 | 0 | 14.09 | 182734 | 6 | 4 | 33 | 150.31 |
| Bacteria | Bacillota_H | GCA-014931565__sp014931565 | 014931565 | 3879629 | 1 | 500 | 1 | 0 | 1 | 13.05 | 182734 | 7 | 4 | 42 | 149.58 |
| Bacteria | Bacteroidota | Fulvivirga__lutea | 017068455 | 4189718 | 1 | 70 | 0 | 0 | 0 | 10.19 | 182734 | 9 | 7 | 21 | 110.45 |
| Bacteria | Bdellovibrionota | Halobacteriovorax_A__vibrionivorans | 003612895 | 3192626 | 1 | 49 | 0 | 0 | 0 | 10.72 | 182734 | 5 | 1 | 21 | 120.02 |
| Bacteria | Bipolaricaulota | Bipolaricaulis__anaerobius | 900465355 | 1340893 | 1 | 500 | 0 | 0 | 1 | 16.66 | 182734 | 6 | 3 | 12 | 115.55 |
| Bacteria | BMS3Abin14 | G020352595__sp020352595 | 020352595 | 2186599 | 1 | 59 | 0 | 0 | 0 | 13.89 | 182734 | 8 | 1 | 10 | 111.42 |
| Bacteria | CAIJMQ01 | CAIJMQ01__sp018812925 | 018812925 | 1488808 | 14 | 324 | 0 | 0 | 0 | 9.68 | 182734 | 7 | 6 | 22 | 117.11 |
| Bacteria | Caldisericota | Caldisericum__exile | 000284335 | 1558104 | 1 | 198 | 0 | 0 | 0 | 10.13 | 182734 | 7 | 5 | 14 | 101.93 |
| Bacteria | Calditrichota | RQOF01__sp020350205 | 020350205 | 4113906 | 1 | 1 | 0 | 0 | 0 | 9.07 | 182734 | 6 | 4 | 30 | 126.45 |
| Bacteria | Calescibacterota | Calescibacterium__sp002898315 | 002898315 | 1899454 | 22 | 0 | 0 | 0 | 0 | 8.13 | 182734 | 6 | 5 | 66 | 117.72 |
| Bacteria | Campylobacterota | Campylobacter_D__jejuni | 001507205 | 1637208 | 1 | 500 | 27 | 6 | 5 | 19.91 | 182734 | 39 | 11 | 28 | 154.24 |
| Bacteria | CG03 | CG03__sp013791745 | 018901505 | 1939027 | 27 | 24 | 0 | 0 | 0 | 9.03 | 182734 | 4 | 2 | 43 | 109.7 |
| Bacteria | CG2-30-53-67 | JAADIO01__sp013151235 | 013151235 | 2653757 | 73 | 500 | 1 | 1 | 0 | 13.38 | 182734 | 6 | 2 | 116 | 143.11 |
| Bacteria | CG2-30-70-394 | CG2-30-70-394__sp001873295 | 002770915 | 2764542 | 149 | 500 | 1 | 0 | 1 | 20 | 182734 | 11 | 3 | 133 | 203.27 |
| Bacteria | Chlamydiota | Chlamydia__trachomatis | 000220105 | 1038314 | 1 | 182 | 0 | 0 | 0 | 12.92 | 182734 | 5 | 3 | 7 | 82.93 |
| Bacteria | Chloroflexota | Dehalococcoides__mccartyi | 000011905 | 1469721 | 1 | 325 | 3 | 3 | 1 | 10.31 | 182734 | 9 | 6 | 10 | 94.8 |
| Bacteria | CLD3 | JABWCP01__sp013360215 | 013360215 | 3766478 | 56 | 20 | 0 | 0 | 0 | 9.3 | 182734 | 11 | 7 | 124 | 140.33 |
| Bacteria | Cloacimonadota | Cloacimonas__acidaminovorans | 000146065 | 2246821 | 1 | 106 | 0 | 0 | 0 | 11.46 | 182734 | 12 | 7 | 12 | 126.44 |
| Bacteria | Coprothermobacte | Coprothermobacter__proteolyticus | 000020945 | 1424913 | 1 | 2 | 1 | 0 | 0 | 8.93 | 182734 | 9 | 4 | 2 | 94.03 |
| Bacteria | CSP1-3 | HRBIN32__sp025060455__X1 | 011053555 | 2474081 | 14 | 500 | 1 | 0 | 1 | 38.73 | 182734 | 12 | 2 | 68 | 147.46 |
| Bacteria | CSSED10-310 | JAFGEQ01__sp016931515 | 016931515 | 2604272 | 24 | 20 | 0 | 0 | 0 | 8.84 | 182734 | 6 | 3 | 36 | 122.92 |

| | | | | | | | | | | | | | | | |
|---|---|---|---|---|---|---|---|---|---|---|---|---|---|---|---|
| Bacteria | Cyanobacteriota | Atelocyanobacterium__thalassa_A | 020885515 | 1510257 | 1 | 500 | 0 | 0 | 0 | 17.09 | 182734 | 9 | 2 | 16 | 101.43 |
| Bacteria | Deferribacterota | Flexistipes__sinusarabici | 000218625 | 2526591 | 1 | 29 | 0 | 0 | 3 | 8.51 | 182734 | 11 | 5 | 20 | 123.51 |
| Bacteria | Deinococcota | G020354625__sp020354625 | 020354625 | 4339659 | 1 | 500 | 0 | 0 | 0 | 11.13 | 182734 | 10 | 6 | 17 | 127.2 |
| Bacteria | Delongbacteria | JACKCN01__sp020634015 | 020634015 | 3933784 | 17 | 500 | 0 | 0 | 1 | 23.22 | 182734 | 11 | 5 | 60 | 148.7 |
| Bacteria | Dependentiae | AalE-18__sp016699045 | 016699045 | 1469267 | 1 | 0 | 0 | 0 | 0 | 7.08 | 182734 | 7 | 1 | 13 | 101.17 |
| Bacteria | Desantisbacteria | UBA1551__sp001871075__X1 | 011389435 | 1980471 | 88 | 0 | 0 | 0 | 0 | 7.49 | 182734 | 5 | 2 | 89 | 113.49 |
| Bacteria | Desulfobacterota | BM004__sp020346205 | 020346205 | 1762565 | 1 | 0 | 0 | 0 | 0 | 8.09 | 182734 | 11 | 6 | 8 | 92.19 |
| Bacteria | Desulfobacterota_ | JAHBUY01__sp019637795 | 019637795 | 6234139 | 12 | 500 | 0 | 0 | 6 | 29.44 | 182734 | 16 | 5 | 83 | 199.61 |
| Bacteria | Desulfobacterota_ | Deferrisoma__camini | 000526155 | 4236552 | 1 | 500 | 1 | 1 | 3 | 25.86 | 182734 | 21 | 7 | 29 | 193.96 |
| Bacteria | Desulfobacterota_ | GCA-014075295__sp014075295 | 011087995 | 980631 | 1 | 0 | 0 | 0 | 0 | 6.93 | 182734 | 8 | 5 | 3 | 102.73 |
| Bacteria | Desulfobacterota_ | CSP1-8__sp020346465 | 020346465 | 2352584 | 1 | 63 | 0 | 0 | 0 | 15.45 | 182734 | 12 | 0 | 13 | 122.91 |
| Bacteria | Desulfobacterota_ | Geobacter__sulfurreducens | 019904315 | 3876396 | 1 | 500 | 2 | 1 | 3 | 18.37 | 182734 | 10 | 3 | 24 | 175.68 |
| Bacteria | Desulfobacterota_ | Syntrophorhabdus__aromaticivorans | 000512235 | 3759485 | 3 | 78 | 0 | 0 | 3 | 9.96 | 182734 | 6 | 4 | 21 | 121.54 |
| Bacteria | Dictyoglomota | Dictyoglomus__thermophilum | 000020965 | 1959988 | 1 | 180 | 0 | 0 | 0 | 11.1 | 182734 | 4 | 3 | 27 | 111.58 |
| Bacteria | Dormibacterota | UBA10449__sp001917815__X1 | 001918245 | 487393 | 21 | 80 | 0 | 0 | 0 | 9.38 | 182734 | 1 | 0 | 5 | 78.51 |
| Bacteria | DRYD01 | DRYD01__sp025059235 | 025059235 | 1406608 | 30 | 500 | 0 | 0 | 0 | 10.92 | 182734 | 7 | 5 | 19 | 97.03 |
| Bacteria | DTU030 | DUMP01__sp012839755 | 019668485 | 2916468 | 1 | 500 | 0 | 0 | 0 | 21.71 | 182734 | 14 | 7 | 14 | 125.82 |
| Bacteria | DUMJ01 | DUMJ01__sp012839925 | 012839925 | 2096858 | 84 | 213 | 0 | 0 | 0 | 9.5 | 182734 | 4 | 2 | 86 | 98.56 |
| Bacteria | Edwardsbacteria | UBA2226__sp001777915 | 001777935 | 2762061 | 8 | 229 | 0 | 0 | 0 | 18.83 | 182734 | 13 | 6 | 29 | 148.37 |
| Bacteria | Eisenbacteria | G020349905__sp020349905 | 020349905 | 2724192 | 1 | 11 | 0 | 0 | 0 | 9.26 | 182734 | 11 | 7 | 14 | 113.49 |
| Bacteria | Elusimicrobiota | Endomicrobium__proavitum | 001027545 | 1588980 | 1 | 21 | 0 | 0 | 0 | 9.68 | 182734 | 9 | 4 | 13 | 102.71 |
| Bacteria | Eremiobacterota | Cybelea__sp019240525 | 019240525 | 2844076 | 5 | 81 | 0 | 0 | 0 | 10.93 | 182734 | 9 | 4 | 20 | 110.01 |
| Bacteria | FCPU426 | Bog-1183__sp003136635 | 003136635 | 3074033 | 61 | 161 | 0 | 0 | 1 | 14.99 | 182734 | 7 | 3 | 115 | 124.32 |
| Bacteria | FEN-1099 | JACKCT01__sp020633895 | 020633915 | 2684210 | 38 | 132 | 0 | 0 | 3 | 16.15 | 182734 | 4 | 2 | 138 | 118.48 |
| Bacteria | Fermentibacterota | UBA6080__sp002436025 | 002436025 | 2860252 | 18 | 0 | 0 | 0 | 0 | 7.55 | 182734 | 9 | 5 | 57 | 124.04 |
| Bacteria | Fibrobacterota | UBA5070__sp016699655 | 016699655 | 5909906 | 1 | 500 | 1 | 0 | 2 | 14.17 | 182734 | 6 | 4 | 26 | 154.98 |
| Bacteria | Firestonebacteria | CAIYPD01__sp903891415 | 903891415 | 2530737 | 86 | 3 | 1 | 1 | 0 | 9.32 | 182734 | 1 | 0 | 139 | 118.67 |
| Bacteria | Fusobacteriota | Fusobacterium__animalis | 019552125 | 2522128 | 1 | 214 | 11 | 4 | 1 | 26.88 | 182734 | 15 | 7 | 52 | 163.28 |
| Bacteria | Gemmatimonadot | G020350785__sp020350785 | 020350785 | 4145679 | 1 | 500 | 0 | 0 | 1 | 15.35 | 182734 | 7 | 4 | 20 | 123.39 |
| Bacteria | Goldbacteria | PGYV01__sp002839855 | 002839855 | 2898434 | 18 | 500 | 0 | 0 | 0 | 12.39 | 182734 | 10 | 5 | 64 | 116.51 |
| Bacteria | Hydrogenedentota | WGMK01__sp016125095 | 016125095 | 5569248 | 44 | 500 | 0 | 0 | 9 | 13.32 | 182734 | 8 | 4 | 218 | 159.34 |
| Bacteria | JAAXHH01 | JAAXHH01__sp022821085 | 022821085 | 3801420 | 7 | 500 | 1 | 0 | 2 | 14.55 | 182734 | 9 | 6 | 33 | 144.57 |
| Bacteria | JAAXVQ01 | JAIORK01__sp021108215 | 021108215 | 3604787 | 67 | 462 | 0 | 0 | 0 | 19.47 | 182734 | 8 | 4 | 141 | 137.52 |
| Bacteria | JABDJQ01 | JAFGGD01__sp016930695 | 016930695 | 3829226 | 60 | 335 | 0 | 0 | 3 | 23.7 | 182734 | 5 | 3 | 197 | 168.93 |
| Bacteria | JAMCPX01 | JAJYLM01__sp021787655 | 021839265 | 1936438 | 14 | 62 | 0 | 0 | 0 | 12.5 | 182734 | 6 | 0 | 34 | 108.04 |
| Bacteria | JdFR-76 | BMS3Abin05__sp002898015 | 913009305 | 3810466 | 86 | 500 | 0 | 0 | 2 | 12.95 | 182734 | 4 | 2 | 216 | 161.58 |
| Bacteria | Krumholzibacterio | G020350885__sp020350885 | 020350885 | 4177611 | 1 | 85 | 0 | 0 | 0 | 9.47 | 182734 | 9 | 2 | 18 | 136.05 |
| Bacteria | KSB1 | JACKDE01__sp020633675 | 020633675 | 5421557 | 8 | 500 | 1 | 0 | 0 | 11.44 | 182734 | 12 | 3 | 31 | 142.61 |
| Bacteria | Margulisbacteria | XYB2-FULL-45-9__sp001771575 | 001771625 | 1668698 | 1 | 384 | 0 | 0 | 0 | 14.68 | 182734 | 10 | 2 | 10 | 110.8 |
| Bacteria | Marinisomatota | G020354025__sp020346745 | 020346745 | 3150876 | 1 | 500 | 0 | 0 | 0 | 9.92 | 182734 | 6 | 6 | 10 | 119.48 |
| Bacteria | Methylomirabilota | Methylomirabilis__oxyfera | 000091165 | 2752855 | 1 | 322 | 1 | 1 | 0 | 14.02 | 182734 | 7 | 2 | 15 | 121.05 |
| Bacteria | Moduliflexota | Moduliflexus__flocculans | 000739515 | 7147165 | 8 | 500 | 1 | 0 | 0 | 10.62 | 182734 | 6 | 2 | 34 | 151.21 |
| Bacteria | Myxococcota | Minicystis__rosea | 001931535 | 16040667 | 1 | 500 | 0 | 0 | 3 | 26.02 | 182734 | 17 | 12 | 89 | 233.89 |
| Bacteria | Myxococcota_A | G020347605__sp020347605 | 020347605 | 3716786 | 1 | 500 | 1 | 0 | 1 | 22.24 | 182734 | 12 | 3 | 36 | 139.18 |
| Bacteria | Nitrospinota | Nitrohelix__vancouverensis | 015698305 | 3351443 | 1 | 500 | 0 | 0 | 3 | 11.59 | 182734 | 12 | 6 | 18 | 125.9 |
| Bacteria | Nitrospirota | Nitrospira_E__moscoviensis | 001273775 | 4589486 | 1 | 500 | 5 | 5 | 1 | 23.9 | 182734 | 15 | 8 | 21 | 138.5 |
| Bacteria | Nitrospirota_A | Leptospirillum_A__rubarum | 001186405 | 2709325 | 1 | 61 | 11 | 11 | 4 | 10.63 | 182734 | 17 | 14 | 17 | 147.91 |
| Bacteria | NPL-UPA2 | NPL-UPA2__sp003574845 | 003574845 | 1008681 | 24 | 26 | 0 | 0 | 0 | 10.36 | 182734 | 6 | 4 | 14 | 87.91 |
| Bacteria | OLB16 | OLB16__sp001567115 | 015075265 | 4640670 | 3 | 133 | 0 | 0 | 1 | 15.68 | 182734 | 9 | 5 | 20 | 144.2 |
| Bacteria | Omnitrophota | Velamenicoccus__archaeovorus | 004102945 | 1998879 | 1 | 500 | 1 | 0 | 0 | 19.28 | 182734 | 9 | 4 | 8 | 102.58 |
| Bacteria | Patescibacteria | UBA12465__sp016432645 | 016432645 | 989238 | 1 | 500 | 1 | 1 | 0 | 11.74 | 182734 | 5 | 3 | 13 | 93.36 |
| Bacteria | Planctomycetota | Pb10988__sp009664345 | 009664345 | 6732599 | 1 | 500 | 1 | 1 | 3 | 10.11 | 182734 | 4 | 2 | 19 | 142.61 |
| Bacteria | Pseudomonadota | Bradyrhizobium__sp015291665 | 024758445 | 8103283 | 1 | 500 | 38 | 27 | 5 | 41.43 | 182734 | 60 | 29 | 51 | 522.84 |
| Bacteria | QNDG01 | JAFGRY01__sp016934955 | 016934955 | 4454378 | 52 | 91 | 0 | 0 | 0 | 9.19 | 182734 | 8 | 3 | 157 | 141.46 |
| Bacteria | Ratteibacteria | DTJG01__sp023818965 | 023818965 | 1399721 | 12 | 152 | 0 | 0 | 0 | 10.93 | 182734 | 4 | 1 | 29 | 106.31 |
| Bacteria | RBG-13-61-14 | RBG-13-61-14__sp001797815 | 001797815 | 4737178 | 132 | 500 | 1 | 1 | 0 | 22.03 | 182734 | 7 | 1 | 248 | 175.74 |
| Bacteria | RBG-13-66-14 | WVWN01__sp020356085 | 020356085 | 2374315 | 1 | 288 | 0 | 0 | 0 | 22.23 | 182734 | 12 | 3 | 11 | 123.7 |
| Bacteria | Riflebacteria | Rifleibacterium__amylolyticum | 009917695 | 5458725 | 35 | 66 | 0 | 0 | 1 | 10.22 | 182734 | 7 | 6 | 85 | 129.75 |
| Bacteria | SAR324 | Arctic96AD-7__sp905182865 | 905182865 | 3267021 | 5 | 500 | 2 | 0 | 0 | 12.04 | 182734 | 13 | 4 | 27 | 123.41 |
| Bacteria | Schekmanbacteria | JACRJM01__sp016219965 | 016219965 | 3619800 | 18 | 50 | 0 | 0 | 0 | 8.73 | 182734 | 3 | 1 | 51 | 125.72 |

| | | | | | | | | | | | | | | | |
|---|---|---|---|---|---|---|---|---|---|---|---|---|---|---|---|
| Bacteria | Spirochaetota | Treponema_B__denticola | 000340705 | 2857527 | 1 | 500 | 4 | 3 | 1 | 15.33 | 182734 | 15 | 9 | 27 | 117.25 |
| Bacteria | Sumerlaeota | Sumerlaea__chitinivorans | 003290465 | 3330219 | 1 | 0 | 0 | 0 | 0 | 8.15 | 182734 | 11 | 5 | 14 | 129.18 |
| Bacteria | Synergistota | Cloacibacillus__evryensis | 000585335 | 3488465 | 1 | 500 | 5 | 3 | 6 | 14.46 | 182734 | 17 | 6 | 18 | 130.5 |
| Bacteria | SZUA-182 | JAJRUS01__sp024277675 | 024277675 | 2712369 | 139 | 492 | 0 | 0 | 0 | 9.98 | 182734 | 2 | 2 | 151 | 119.15 |
| Bacteria | SZUA-79 | Acididesulfobacter__guangdongensis | 004195045 | 2244839 | 5 | 500 | 0 | 0 | 1 | 17.1 | 182734 | 14 | 7 | 43 | 166.07 |
| Bacteria | TA06 | SM1-40__sp001303705 | 001302835 | 2323841 | 121 | 63 | 0 | 0 | 0 | 7.97 | 182734 | 3 | 3 | 85 | 119.97 |
| Bacteria | Tectomicrobia | SXND01__sp005777235 | 945893115 | 3278621 | 125 | 500 | 0 | 0 | 2 | 9.67 | 182734 | 4 | 3 | 120 | 117.72 |
| Bacteria | Thermodesulfobiot | Thermodesulfobium__acidiphilum | 003057965 | 1796979 | 1 | 202 | 0 | 0 | 0 | 10.42 | 182734 | 10 | 4 | 20 | 129.21 |
| Bacteria | Thermosulfidibacte | Thermosulfidibacter__takaii | 001547735 | 1816671 | 1 | 40 | 0 | 0 | 1 | 8.67 | 182734 | 8 | 4 | 7 | 105.5 |
| Bacteria | Thermotogota | Thermotoga__maritima | 000978535 | 1859583 | 1 | 500 | 0 | 0 | 0 | 12.17 | 182734 | 10 | 4 | 8 | 99.84 |
| Bacteria | UBA10199 | GCA-016699445__sp016699445 | 016699445 | 2780885 | 1 | 9 | 0 | 0 | 2 | 8.54 | 182734 | 9 | 6 | 15 | 113.04 |
| Bacteria | UBA3054 | UBA3054__sp017557885 | 017557885 | 2269068 | 13 | 500 | 1 | 1 | 1 | 22.37 | 182734 | 15 | 6 | 36 | 134.76 |
| Bacteria | UBA6262 | Fen-1174__sp003153935 | 003153935 | 1769019 | 36 | 22 | 0 | 0 | 0 | 9.06 | 182734 | 2 | 2 | 46 | 107.07 |
| Bacteria | UBA6266 | UBA6266__sp023228325 | 023228325 | 2286510 | 39 | 50 | 0 | 0 | 1 | 9.43 | 182734 | 8 | 5 | 20 | 142.16 |
| Bacteria | UBA8248 | Bin107__sp022828125 | 022828125 | 3967823 | 32 | 500 | 1 | 1 | 1 | 11.59 | 182734 | 12 | 5 | 120 | 170.2 |
| Bacteria | UBA9089 | JAHJRX01__sp018830565 | 018818825 | 1952654 | 80 | 207 | 0 | 0 | 0 | 12.7 | 182734 | 1 | 1 | 133 | 113.93 |
| Bacteria | UBP14 | UBA6098__sp002428525 | 002428525 | 2534796 | 35 | 114 | 0 | 0 | 0 | 8.44 | 182734 | 6 | 3 | 88 | 129.75 |
| Bacteria | UBP15 | UBA6099__sp002435745 | 002435745 | 2590492 | 62 | 170 | 1 | 1 | 0 | 11.06 | 182734 | 4 | 1 | 79 | 113.5 |
| Bacteria | UBP17 | GCA-2402105__sp023954715 | 023954715 | 4638379 | 38 | 39 | 0 | 0 | 0 | 8.47 | 182734 | 6 | 4 | 159 | 126.28 |
| Bacteria | UBP18 | UBA7526__sp002478245 | 002478245 | 1174104 | 90 | 0 | 0 | 0 | 0 | 7.03 | 182734 | 4 | 1 | 20 | 87.27 |
| Bacteria | UBP6 | UBA1177__sp902774115 | 902774115 | 2544559 | 15 | 92 | 1 | 0 | 0 | 11.27 | 182734 | 8 | 2 | 31 | 130.13 |
| Bacteria | UBP7 | UBA6624__sp002716355 | 002716355 | 1280514 | 16 | 500 | 0 | 0 | 1 | 9.48 | 182734 | 8 | 4 | 45 | 108.98 |
| Bacteria | Verrucomicrobiota | Akkermansia__muciniphila_B | 018846995 | 3202594 | 1 | 224 | 18 | 18 | 4 | 15.06 | 182734 | 25 | 21 | 20 | 150.78 |
| Bacteria | Wallbacteria | UBA9980__sp002069765 | 022703765 | 5042914 | 209 | 17 | 0 | 0 | 1 | 9.55 | 182734 | 1 | 1 | 334 | 148.76 |
| Bacteria | WOR-3 | SM23-42__sp020353055 | 020353055 | 2721997 | 1 | 250 | 0 | 0 | 0 | 11.03 | 182734 | 3 | 0 | 11 | 99.51 |
| Bacteria | Zixibacteria | JAFGAL01__sp020343475 | 020343475 | 2882114 | 1 | 234 | 1 | 1 | 0 | 9.95 | 182734 | 4 | 2 | 14 | 122.81 |

| | | | | *Taxonomic* | | | | | *Search* | | | | |
|---|---|---|---|---|---|---|---|---|---|---|---|---|---|
| **GTDB species** | **GCA** | **Genome length (bp)** | **No. contigs** | **No. attBs** | **attB candidates** | **attBs in tRNA genes** | **Occupied** | **Runtime (s)** | **No. attBs** | **attB candidates** | **attBs in tRNA genes** | **Occupied** | **Runtime (s)** |
| Acidithiobacillus_ferrooxidans | 000021485 | 2982398 | 1 | 293 | 24 | 18 | 2 | 39.32 | 182734 | 29 | 15 | 18 | 270.51 |
| Acinetobacter_baumannii | 009759685 | 4040269 | 2 | 500 | 34 | 10 | 7 | 58.37 | 182734 | 49 | 17 | 33 | 257.92 |
| Bacillus_coagulans_DSM2314 | 006716385 | 3674010 | 1 | 482 | 4 | 2 | 0 | 48.92 | 182734 | 15 | 9 | 25 | 153.84 |
| Bacillus_coagulans_MA-13 | 004359975 | 2979001 | 116 | 500 | 3 | 2 | 0 | 46.89 | 182734 | 6 | 3 | 177 | 148.31 |
| Bacillus_licheniformis_DSM13 | 000011645 | 4222598 | 1 | 500 | 24 | 9 | 2 | 39.6 | 182734 | 33 | 13 | 21 | 163.61 |
| Bacillus_subtilis | 000009045 | 4215607 | 1 | 500 | 17 | 2 | 2 | 38.62 | 182734 | 27 | 8 | 22 | 151.76 |
| Bacteroides_fragilis | 000025985 | 5241702 | 2 | 500 | 38 | 12 | 9 | 64.08 | 182734 | 58 | 24 | 26 | 294.56 |
| Clostridioides_difficile_DSM28196 | 003482035 | 4257933 | 1 | 500 | 29 | 0 | 25 | 74.44 | 182734 | 48 | 5 | 61 | 217.44 |
| Clostridium_acetobutylicum_ATCC824 | 000008765 | 4132882 | 2 | 500 | 0 | 0 | 1 | 34.19 | 182734 | 11 | 3 | 44 | 152.78 |
| Clostridium_ljungdahlii_DSM13528 | 000143685 | 4630066 | 1 | 214 | 1 | 0 | 4 | 30.42 | 182734 | 6 | 3 | 52 | 154.12 |
| Clostridium_tyrobutyricum_ATCC25755 | 000359585 | 3010772 | 76 | 286 | 8 | 3 | 3 | 36.23 | 182734 | 6 | 3 | 296 | 139.31 |
| Corynebacterium_diphtheriae | 001457455 | 2463667 | 1 | 203 | 16 | 7 | 1 | 33.26 | 182734 | 29 | 13 | 8 | 120.31 |
| Corynebacterium_glutamicum_ATCC130 | 000011325 | 3309402 | 1 | 152 | 10 | 7 | 2 | 34.13 | 182734 | 22 | 11 | 15 | 115.48 |
| Cupriavidus_necator_H16 | 004798725 | 7507244 | 3 | 500 | 27 | 17 | 5 | 61.89 | 182734 | 47 | 16 | 59 | 779.98 |
| Enterococcus_faecalis | 000392875 | 2881403 | 3 | 500 | 38 | 5 | 9 | 53.55 | 182734 | 42 | 6 | 28 | 172.22 |
| Escherichia_coli | 003697165 | 5097770 | 2 | 500 | 44 | 8 | 7 | 69.06 | 182734 | 157 | 26 | 27 | 3825.99 |
| Francisella_tularensis | 000008985 | 1892776 | 1 | 500 | 3 | 2 | 0 | 32.51 | 182734 | 18 | 8 | 19 | 139.32 |
| Haloferax_volcanii | 000025685 | 4012905 | 5 | 500 | 10 | 5 | 8 | 64.14 | 182734 | 12 | 8 | 38 | 129.66 |
| Helicobacter_pylori | 900478295 | 1701949 | 1 | 500 | 22 | 1 | 0 | 37.57 | 182734 | 33 | 5 | 8 | 107.89 |
| Klebsiella_pneumoniae | 000742135 | 5545789 | 5 | 500 | 37 | 9 | 9 | 69.01 | 182734 | 100 | 28 | 36 | 2585.67 |
| Leptospirillum_ferriphilum | 000755505 | 2405900 | 18 | 67 | 12 | 12 | 2 | 23.99 | 182734 | 18 | 13 | 52 | 131.56 |
| Listeria_monocytogenes | 900187225 | 2900472 | 1 | 500 | 38 | 7 | 0 | 45.45 | 182734 | 50 | 9 | 19 | 149.07 |
| Methylobacterium_aquaticum | 001043915 | 7518394 | 512 | 500 | 7 | 7 | 2 | 49.33 | 182734 | 2 | 2 | 364 | 312.2 |
| Methylorubrum_extorquens | 900234795 | 5786956 | 1 | 500 | 23 | 20 | 5 | 63.07 | 182734 | 39 | 20 | 43 | 324.83 |
| Mycobacterium_tuberculosis | 000195955 | 4411533 | 1 | 500 | 24 | 16 | 4 | 44.12 | 182734 | 31 | 17 | 35 | 242.92 |
| Neisseria_gonorrhoeae | 003315235 | 2169591 | 72 | 500 | 15 | 10 | 8 | 43.93 | 182734 | 18 | 9 | 85 | 178.68 |
| Novosphingobium_aromaticivorans_DSM | 000013325 | 4233317 | 3 | 500 | 10 | 8 | 3 | 43.15 | 182734 | 31 | 19 | 35 | 294.17 |
| Pseudomonas_aeruginosa | 001457615 | 6316980 | 1 | 500 | 46 | 15 | 5 | 73.67 | 182734 | 91 | 24 | 41 | 1286.36 |
| Pseudomonas_fluorescens_SBW25 | 000009225 | 6722540 | 1 | 500 | 29 | 16 | 8 | 59.5 | 182734 | 59 | 23 | 39 | 806.04 |
| Pseudomonas_putida_KT2440 | 000007565 | 6181874 | 1 | 500 | 34 | 14 | 9 | 62.31 | 182734 | 62 | 20 | 33 | 636.08 |
| Pseudomonas_putida_S12 | 000495455 | 6382436 | 2 | 500 | 37 | 15 | 6 | 69.06 | 182734 | 67 | 18 | 35 | 701.14 |
| Rhodobacter_sphaeroides_ATCC17023- | 000012905 | 4602984 | 7 | 500 | 9 | 6 | 10 | 51.43 | 182734 | 33 | 18 | 40 | 256.18 |
| Salmonella_enterica | 000006945 | 4951385 | 2 | 500 | 36 | 10 | 7 | 71.15 | 182734 | 139 | 24 | 24 | 4556.94 |
| Staphylococcus_aureus | 001027105 | 2782564 | 2 | 500 | 25 | 2 | 4 | 50.62 | 182734 | 33 | 3 | 30 | 142.77 |
| Streptococcus_pneumoniae | 001457635 | 2110969 | 1 | 500 | 44 | 3 | 16 | 56.34 | 182734 | 94 | 5 | 21 | 288.67 |
| Streptomyces_griseus_ATCC13273 | 003610995 | 7328672 | 1 | 500 | 32 | 17 | 5 | 72.38 | 182734 | 60 | 20 | 54 | 602.05 |
| Streptomyces_venezualae | 008639165 | 8326299 | 1 | 500 | 22 | 11 | 12 | 62.7 | 182734 | 61 | 23 | 55 | 567.01 |
| Sulfolobus_acidocaldarius | 000012285 | 2225960 | 1 | 33 | 0 | 0 | 0 | 24.15 | 182734 | 1 | 0 | 13 | 82.34 |
| Synechococcus_elongatus | 000817325 | 2744629 | 3 | 72 | 1 | 1 | 0 | 27.36 | 182734 | 14 | 10 | 7 | 96.73 |
| Vibrio_cholerae | 000621645 | 4024579 | 62 | 500 | 26 | 9 | 3 | 43.43 | 182734 | 33 | 11 | 80 | 174.98 |
| Zymomonas_mobilis_ZM4 | 003054575 | 2227818 | 5 | 111 | 0 | 0 | 0 | 25.09 | 182734 | 14 | 10 | 13 | 194.24 |
