## Supplementary Table 2 for "Integrase-On-Demand: Bioprospecting integrases for targeted genomic insertion of genetic cargo"

### Pseudomonas putida S12

| query_contig | strand | query_coord | query_attB | integrase | isleID | ref_scaffold/coords | source | support | isle_type |
|---|---|---|---|---|---|---|---|---|---|
| CP009974.1 | minus | 3663-3595 | GGTTATGCTCCGGA | Y-P2Arm_02846 | 003936165.25.C | RHRX01000032.1/11092-1+RHRX01000011.1/134460-120897 | TIGER | 80 | other |
| CP009974.1 | minus | 4863-4829 | AGGAGTCAACGTGG | Y-Tn916Arm_01138 | 900187555.6.sohB | FYDW01000005.1/174620-168219 | TIGER | 182 | Phage2 |
| CP009974.1 | minus | 56722-56652 | GTTTAAGGCAACGG | Y-P2Arm_03837 | 903684265.4.S | CAHPQU010000018.1/102094-104191+CAHPQU010000057.1/1-1469 | TIGER | 25 | other |
| CP009974.1 | minus | 141734-141697 | GTTCGAGTCTCGTTT | Y-SXTArm_05826 | 021602445.34.G | BQII01000065.1/10464-27298+BQII01000065.1/10464-27298 | TIGER | 25 | other |
| CP009974.1 | plus | 259317-259365 | GAACTGCGTGCACT | Y-SXTArm_04049 | 023461295.42.ETF_QO | JAEYMN010000182.1/27573-1+JAEYMN010000025.1/59637-45654 | TIGER | 7 | other |
| CP009974.1 | minus | 364171-364080 | TTACCCGTTAACCTG | Y-Tn916Arm_05122 | 021283545.11.Archaea_SRP | JAJSRM010000045.1/4558-1+JAJSRM010000069.1/22326-15821 | TIGER | 63 | other |
| CP009974.1 | minus | 524540-524477 | TTGGATAGAGCATCC | Y-SXTArm_02325 | 016008995.9.R | JADUCG010000015.1/29175-38513 | Islander,TIGE | 59 | other |
| CP009974.1 | plus | 641798-641838 | GCGTGACGAGCCA | S-Int_02028 | 016907075.7.DUF748\|AAA | JAFBBH010000001.1/2215329-2222503 | TIGER | 23 | other |
| CP009974.1 | minus | 688697-688650 | ACAGTACCTAGGGT | Y-P2Arm_05871 | 000498395.4.S | CP010979.1/6478889-6483051 | TIGER | 84 | other |
| CP009974.1 | minus | 706350-706310 | GCCAGTCGCGGGT | S-Int_04377 | 002906815.7.tol-pal_system_protein_YbgF\|queE | MING01000083.1/1634708-1627560 | TIGER | 196 | other |
| CP009974.1 | plus | 813670-813704 | TCAATGCCAGGTTCT | Y-Tn916Arm_03719 | 021283195.6.HYP | JAJSRH010000033.1/34795-28475 | TIGER | 193 | Phage2 |
| CP009974.1 | plus | 845618-845652 | TTTCAAGAACCGGAC | Y-SXTArm_01088 | 002953115.12.bfd\|bfr | CP026332.1/1228357-1216200 | TIGER | 195 | other |
| CP009974.1 | plus | 1121771-1121805 | GTTCGAGTCTCCCT | Y-P2Arm_04621 | 001653615.34.L | LSUZ01000199.1/41005-6509 | Islander,TIGE | 26 | Phage1 |
| CP009974.1 | plus | 1261821-1261849 | CTTCCGCTTCAACG | Y-SXTArm_03220 | 004614155.26.ychF | SPUU01000036.1/47681-53714+SPUU01000035.1/55064-35118 | TIGER | 46 | other |
| CP009974.1 | minus | 1360887-1360846 | GGTAGAGCAGCGC | Y-P2Arm_05361 | 021283035.11.T | JAJSQQ010000063.1/11491-213 | Islander | 187 | Phage1 |
| CP009974.1 | plus | 1478260-1478293 | CCCCGAGAAGCCG | Y-SXTArm_00656 | 002356095.67.prtR\|TonB_dep_Rec | AP015029.1/709926-642683 | TIGER | 37 | other |
| CP009974.1 | plus | 1518981-1519005 | GACTGGGGGCGCG | Y-TnpA_01692 | 004519745.102.alsT | PZKR01000021.1/102440-177533+PZKR01000008.1/82264-55792 | TIGER | 12 | other |
| CP009974.1 | plus | 1637681-1637745 | GAGCAGAGGACTCA | Y-P2Arm_04153 | 000498395.10.J | CP010979.1/4574849-4584722 | Islander,TIGE | 83 | other |
| CP009974.1 | plus | 1859950-1860016 | GAGTACTGCCCTCC | Y-P2Arm_00208 | 014217705.9.R | CP046872.1/251211-241839 | Islander,TIGE | 83 | Phage2 |
| CP009974.1 | minus | 2306556-2306510 | CACAGGCGTAAAAA | Y-SXTArm_04117 | 001753955.5.yigB\|HYP | BCBP01000050.1/23473-18452 | TIGER | 118 | ICE1 |
| CP009974.1 | minus | 2984352-2984303 | AGTACTGGCGGACG | Y-BPP1Arm_04851 | 000271965.40.Z | CP015876.1/5210942-5250776 | Islander,TIGE | 97 | Phage1 |
| CP009974.1 | minus | 3332631-3332593 | ATGAAATCGACGAGT | Y-Tn916Arm_03030 | 019219145.7.nadB | JAHUWI010000003.1/14545-21112 | TIGER | 190 | other |
| CP009974.1 | minus | 3496657-3496620 | GAAATCAACCAGTTC | Y-Tn916Arm_05199 | 013386405.7.nadB | JACARV010000075.1/48544-55191 | TIGER | 39 | other |
| CP009974.1 | minus | 3624898-3624860 | GTAAGGACTTATGTG | Y-BPP1Arm_00722 | 021282585.32.asnB | JAJSPR010000024.1/13977-1+JAJSPR010000015.1/121491-103117 | TIGER | 109 | Phage1 |
| CP009974.1 | plus | 3741708-3741759 | AATCGTATCAAGGGT | Y-BPP1Arm_01300 | 000764405.50.S | JOJY01000001.1/1286654-1336311 | Islander,TIGE | 96 | Phage1 |
| CP009974.1 | plus | 3771933-3771972 | TCGCGAGTTCGAGT | Y-SXTArm_01796 | 000367825.94.G | APBQ01000056.1/93015-1+APBQ01000112.1/1-590 | TIGER | 66 | ICE2 |
| CP009974.1 | minus | 4015655-4015617 | AATCCGCACCCCG | S-Int_01475 | 001312165.12.bvgS | BBDB01000027.1/13048-761 | TIGER | 184 | other |
| CP009974.1 | minus | 4037268-4037235 | GTTCGAATCCTGTCT | Y-SXTArm_05164 | 002165665.9.P | NHBC01000017.1/401597-410151 | Islander,TIGE | 119 | other |
| CP009974.1 | minus | 4091592-4091553 | CTTCAGCGTTGCCC | Y-P2Arm_03735 | 003671955.52.dusC | CP022561.1/4044796-4096450 | TIGER | 19 | Phage1 |
| CP009974.1 | plus | 4202755-4202829 | AGACACACTGGATTT | Y-P2Arm_03960 | 001320655.39.L | BCBK01000084.1/2994-1+BCBK01000083.1/104792-68316 | TIGER | 98 | Phage1 |
| CP009974.1 | minus | 4250453-4250417 | TTATGACAGCGTGGA | Y-SXTArm_00202 | 000367825.52.queD | APBQ01000005.1/47698-1+APBQ01000125.1/1-4693 | TIGER | 67 | Phage1 |
| CP009974.1 | plus | 4370757-4370830 | TTGGTTAGAGCACCA | Y-P2Arm_01214 | 002835905.40.V | PJCG01000008.1/69296-29786 | Islander,TIGE | 16 | Phage1 |
| CP009974.1 | minus | 5224650-5224624 | CGGCAACGTTCA-A | Y-SXTArm_01372 | 021282465.58.HYP\|HYP | JAJSRF010000001.1/131429-73346 | TIGER | 107 | ICE2 |
| CP009974.1 | plus | 5251460-5251486 | CTGCTTCATGCCGG | Y-Cyan_03156 | 000007565.17.UPF0157 | AE015451.2/3383569-3366551 | TIGER | 29 | other |
| CP009974.1 | plus | 5288587-5288624 | TTGATCTCCACCGC | Y-BPP1Arm_00929 | 016845885.57.yheS | CP069317.1/948766-1006083 | TIGER | 14 | Phage1 |
| CP009974.1 | plus | 5453558-5453601 | CCAGATGCGCGTGC | Y-SXTArm_01106 | 021282705.119.azr | JAJSQG010000019.1/43103-1+JAJSQG010000003.1/181800-106197 | TIGER | 67 | Phage1 |
| CP009974.1 | plus | 5776009-5776041 | ATACACAGCAGACTT | Y-SXTArm_02201 | 000987155.62.HYP\|yphA | AYRY01000007.1/154563-92805 | TIGER | 142 | other |

**Occupied Integrases Tested**

| query_contig | strand | query_coord | query_attB | integrase | isleID | ref_scaffold/coords | source | support | isle_type | |
|---|---|---|---|---|---|---|---|---|---|---|
| CP009974.1 | minus | 5199833-5199794 | GTTCTGGGTGGCGG | S-Int_05038 | 000264665.40.eamB | CP003588.1/5514732-5555071 | TIGER | 147 | Phage1 | NOTE: flagged as questionable, has ID block in genome 26-times |
| CP009974.1 | plus | 4394684-4394711 | GCGCACTTGCAT-G | Y-SXTArm_03422 | 000271965.63.P | CP015876.1/3754094-3690895 | Islander,TIGE | 56 | other | NOTE: flagged as questionable, attB overlaps a predicted island (CP009974.1/4394681-4412437) |

### Pseudomonas putida KT2440

| query_contig | strand | query_coord | query_attB | integrase | isleID | ref_scaffold/coords | source | support | isle_type |
|---|---|---|---|---|---|---|---|---|---|
| AE015451.2 | minus | 255336-255276 | ATAGAGTACTGCCC | Y-P2Arm_05397 | 001941965.11.R | MKZO01000058.1/294596-283401 | Islander | 21 | Phage1 |
| AE015451.2 | minus | 335029-334963 | TAGAGCAACTGACTT | Y-P2Arm_05338 | 002906815.31.T | MING01000083.1/2646824-2677956 | Islander,TIGE | 65 | Phage1 |
| **AE015451.2** | **minus** | **467174-467110** | **GAGCAGAGGACTC** | **Y-SXTArm_06036** | **021282465.12.M** | **JAJSRF010000008.1/77721-65525** | **Islander,TIGE** | **83** | **Phage2** |
| AE015451.2 | minus | 585762-585738 | GACTGGGGGCGCG | Y-TnpA_01692 | 004519745.102.alsT | PZKR01000021.1/102440-177533+PZKR01000008.1/82264-55792 | TIGER | 12 | other |
| **AE015451.2** | **minus** | **624008-623967** | **CCCCGAGAAGCCG** | **Y-SXTArm_02762** | **001319995.25.prtR\|Hemerythrin** | **BCAS01000034.1/6348-30848** | **TIGER** | **41** | **other** |
| AE015451.2 | plus | 741342-741383 | GGTAGAGCAGCGC | Y-P2Arm_05361 | 021283035.11.T | JAJSQQ010000063.1/11491-213 | Islander | 187 | Phage1 |
| AE015451.2 | minus | 836273-836246 | CTTCCGCTTTAACGT | Y-TnpA_05248 | 002094775.30.ychF | NBWA01000007.1/8360-1+NBWA01000045.1/1-21439 | TIGER | 73 | other |
| AE015451.2 | minus | 968921-968887 | GTTCGAGTCTCCCT | Y-P2Arm_04621 | 001653615.34.L | LSUZ01000199.1/41005-6509 | Islander,TIGE | 26 | Phage1 |
| AE015451.2 | minus | 1242657-1242623 | TTTCAAGAACCGGAC | Y-SXTArm_01088 | 002953115.12.bfd\|bfr | CP026332.1/1228357-1216200 | TIGER | 195 | other |
| AE015451.2 | minus | 1283461-1283427 | TCAATGCCAGGTTCT | Y-Tn916Arm_03719 | 021283195.6.HYP | JAJSRH010000033.1/34795-28475 | TIGER | 193 | Phage2 |
| AE015451.2 | plus | 1400871-1400911 | GCCAGTCGCGGGT | S-Int_04377 | 002906815.7.tol-pal_system_protein_YbgF\|queE | MING01000083.1/1634708-1627560 | TIGER | 196 | other |
| AE015451.2 | plus | 1402427-1402492 | GAGCAGTTGGCTTTT | Y-KX_02112 | 001753975.34.K | BCBR01000027.1/26113-48345+BCBR01000115.1/19369-7911 | TIGER | 158 | ICE2 |
| AE015451.2 | plus | 1418398-1418445 | ACAGTACCTAGGGT | Y-BPP1Arm_05220 | 021602445.41.S | BQII01000046.1/36679-1+BQII01000082.1/1-4003 | TIGER | 84 | Phage1 |
| AE015451.2 | minus | 1625424-1625386 | ATGAAATCGACGAGT | Y-Tn916Arm_03030 | 019219145.7.nadB | JAHUWI010000003.1/14545-21112 | TIGER | 190 | other |
| AE015451.2 | minus | 1829324-1829287 | GAAATCAACCAGTTC | Y-Tn916Arm_05199 | 013386405.7.nadB | JACARV010000075.1/48544-55191 | TIGER | 39 | other |
| **AE015451.2** | **minus** | **1951455-1951417** | **GTAAGGACTTATGTG** | **Y-BPP1Arm_03980** | **000226035.38.asnB** | **CP015202.1/4197550-4235989** | **TIGER** | **109** | **Phage1** |
| AE015451.2 | plus | 2069117-2069168 | AATCGTATCAAGGGT | Y-BPP1Arm_01300 | 000764405.50.S | JOJY01000001.1/1286654-1336311 | Islander,TIGE | 96 | Phage1 |
| AE015451.2 | plus | 2103013-2103050 | GCGAGTTCGAGTCT | Y-SXTArm_03109 | 005223245.111.G | SWEL01000003.1/919519-808116 | TIGER | 32 | ICE2 |
| AE015451.2 | minus | 2420647-2420614 | GTTCGAATCCTGTCT | Y-SXTArm_05164 | 002165665.9.P | NHBC01000017.1/401597-410151 | Islander,TIGE | 119 | other |
| AE015451.2 | plus | 2682128-2682153 | GACCCGCTGTACAT | Y-TnpA_02250 | 021283035.10.acsA | JAJSQQ010000016.1/3182-1+JAJSQQ010000024.1/1-6372 | TIGER | 169 | other |
| AE015451.2 | minus | 3105111-3105068 | CCAGATGCGCGTGC | Y-SXTArm_01106 | 021282705.119.azr | JAJSQG010000019.1/43103-1+JAJSQG010000003.1/181800-106197 | TIGER | 67 | Phage1 |
| AE015451.2 | plus | 3233802-3233839 | GACCCATTCCAGCT | Y-BPP1Arm_00923 | 002810225.55.gbpR | PIJT01000013.1/172040-116961 | TIGER | 167 | Phage1 |
| AE015451.2 | minus | 3340828-3340790 | CTTGATTTCTACCGC | Y-BPP1Arm_04780 | 002948105.12.yheS | CP026675.1/5304046-5291816 | TIGER | 87 | other |
| AE015451.2 | plus | 4361701-4361733 | ATACACAGCAGACTT | Y-SXTArm_02201 | 000987155.62.HYP\|yphA | AYRY01000007.1/154563-92805 | TIGER | 142 | other |
| AE015451.2 | minus | 4428775-4428741 | AGGAGTCAACGTGG | Y-Tn916Arm_01138 | 900187555.6.sohB | FYDW01000005.1/174620-168219 | TIGER | 182 | Phage2 |
| AE015451.2 | minus | 4632584-4632523 | GTCGCGAGTTCGAG | Y-SXTArm_03827 | 001655295.17.G | CP011525.1/4116293-4099223 | Islander,TIGE | 66 | other |
| AE015451.2 | plus | 4749877-4749925 | GAACTGCGTGCACT | Y-SXTArm_04049 | 023461295.42.ETF_QO | JAEYMN010000182.1/27573-1+JAEYMN010000025.1/59637-45654 | TIGER | 7 | other |
| AE015451.2 | minus | 4858474-4858377 | TTACCCGTTAACCTG | Y-Tn916Arm_05122 | 021283545.11.Archaea_SRP | JAJSRM010000045.1/4558-1+JAJSRM010000069.1/22326-15821 | TIGER | 63 | other |
| AE015451.2 | plus | 5192624-5192664 | GCGTGACGAGCCA | S-Int_02028 | 016907075.7.DUF748\|AAA | JAFBBH010000001.1/2215329-2222503 | TIGER | 23 | other |
| AE015451.2 | plus | 5390394-5390422 | CCCCCCGGCTCCA | Y-SXTArm_03048 | 943913085.14.Z | CALTWL010000015.1/1102-15208 | TIGER | 69 | other |
| AE015451.2 | plus | 5475364-5475407 | CCGGGCAACGCGG | S-Int_05142 | 024971855.8.Pur_DNA_glyco | CP079827.1/5582786-5590516 | TIGER | 113 | ICE1 |
| AE015451.2 | plus | 5965550-5965596 | CACAGGCGTAAAAA | Y-SXTArm_04117 | 001753955.5.yigB\|HYP | BCBP01000050.1/23473-18452 | TIGER | 118 | ICE1 |

### Synechococcus elongatus UTEX 2973

| query_con | strand | query_coord | query_attB | integrase | isleID | ref_scaffold/coords | source | support | isle_type |
|---|---|---|---|---|---|---|---|---|---|
| CP006471. | minus | 13106-13040 | GTGGTAAAACCTTAG | Y-Brujita_00620 | 003957805.84.G | CP033061.1/601056-516596 | Islander,TIGE | 10 | Phage1 |
